# Crosstalk of Histone Modifications in the Healthy Human Immune System

**DOI:** 10.1101/2022.01.21.477300

**Authors:** Denis Dermadi, Laurynas Kalesinskas, Ananthakrishnan Ganesan, Alex Kuo, Peggie Cheung, Sarah Cheng, Mei Dvorak, Thomas J. Scriba, Aida Habtezion, Michele Donato, Paul J. Utz, Purvesh Khatri

## Abstract

Chromatin remodeling through post-translational modifications of histone tails (HPTM) is fundamental for regulation and maintenance of DNA-centered processes. Systems level understanding of coordination and interactions between HPTMs and their impact on the functional state of the immune cells remain unexplored due to the technical reasons. We leveraged large biologically heterogeneous data (>27 million cells), comprising of primary human immune cells profiled for 33 HPTMs and 4 histone variants at the single-cell level using high-dimensional mass cytometry (EpiTOF), to discover and map relations between HPTMs at the systems level. Briefly, we elucidated a comprehensive epigenetic network of HPTM interactions, discovered a novel subset of hematopoietic progenitors with distinct epigenetic profile, and revealed hitherto undescribed associations between a decrease in global methylations, modulation of one-carbon metabolism, and immune cell life span. Ultimately our work lays a foundation for future studies aimed at understanding complexity of HPTM interactions in immune response in infectious or autoimmune diseases, cancers, and vaccination.

---

Eukaryotic genetic material is organized into chromatin, a highly structured polymer of DNA and histones H3, H4, H2A, and H2B^1^. Histones feature flanking N-terminal tails that are post-translationally modified at lysines, arginines, and serines. Histone posttranslational modifications (HPTMs) are largely associated with relaxation or compaction of chromatin; therefore, directly impacting regulation, activation, and repression of transcription. Additionally, HPTMs regulate a variety of distinct biological functions and cell states such as DNA damage and repair signaling, maintenance of cell cycle progression, quiescence, and apoptosis. Recent studies have demonstrated complex and nuanced crosstalk between multiple HPTMs, referred to as “histone language”^2^, as the basis of transcriptional regulation in response to the environment. Histone language is based on spatial co-localization and the simultaneous presence of 2 or more PTMs on (1) a single histone within a nucleosome (*cis* histone pathway) and/or (2) PTMs on different histones within the same or adjacent nucleosome (*trans* histone pathway)^3,4^. For example, phosphorylation of serine 10 (S10) on H3 (H3S10ph) promotes the acetylation of H3K14 (H3K14ac) and inhibits methylation of H3K9 (H3K9me) on the same histone tail (*cis* pathway)^5,6^. In contrast, ubiquitination of H2B modulates multiple methylation events on H3 (*trans* pathway)^7–9^.

To date, the histone language has been mostly studied *in vitro* or in lower eukaryotes using direct mutagenesis of sites of histone modifications. Similar experimental designs are very difficult, if not impossible, *in vivo* and in higher eukaryotes, including humans^4^. Discovery of the histone language has been driven largely by mass spectrometry and ChIP-seq, both gold standards in detecting novel HPTMs and assessing the spatial distribution of modifications across the genome, respectively. However, although unrivaled in revealing spatial and functional roles of PTMs, ChIP-seq is limited to the measurement of 2-3 histone modifications simultaneously, requires large quantities of a sample, and is labor-intensive^10^. Despite the feasibility of ChIP-seq at a single-cell resolution, it is still limited to a few thousand single cells and simultaneous detection of at most two HPTMs^11,12^.

Here, we present the first single-cell resolution study of the nuanced crosstalk between HPTMs in the healthy human immune system using EpiTOF^13^, a high-throughput mass cytometry technology. We profiled 37 HPTMs and histone variants along with 16 cellular phenotypic markers across almost 28 million peripheral blood mononuclear cells (PBMCs) from 5 independent cohorts comprising 83 healthy individuals. The wealth of data at a singlecell resolution presents an unprecedented opportunity to investigate correlations between HPTMs, and the resulting regulatory network, which can potentially reveal novel HPTM interactions in the histone language. Using these data, we (1) elucidated a comprehensive epigenetic network of HPTM interactions in healthy human PBMCs, (2) discovered a previously unreported subset of hematopoietic progenitors with distinct epigenetic profile, and (3) revealed hitherto undescribed associations between a decrease in global cellular methylations, metabolism modulation, and immune cell life span.

## Results

### Data collection

We profiled 27,841,803 PBMCs from 83 healthy individuals across 5 independent cohorts matched for age and sex using EpiTOF. These cohorts included subjects from two continents with ages ranging from 16 years to 80 years old (**Table S1**). We analyzed three previously published cohorts from the US (BR1, BR2, Twins)^13^, a cohort of healthy adults from the US (Stanford) and a cohort of adolescents from South Africa (South Africa). We profiled 37 HPTMs and histone variants divided into two panels: one predominantly measures histone methylations (methylation panel), and the other histone acetylations, phosphorylations, and ubiquitinations (acetylation panel). Both panels include 16 phenotypic cell surface markers for immune cells of human peripheral blood^13^. We determined correlations between HPTMs abundances in each major immune cell type in the PBMCs from healthy individuals, including hematopoietic progenitor cells (HPCs), plasmacytoid dendritic cells (pDCs), myeloid dendritic cells (mDCs), natural killer (NK) cells, NKT, B, CD4, and CD8 T cells, classical monocytes (cMOs), intermediate monocytes (iMOs) and non-classical monocytes (ncMOs) (**Methods**).

### Discovery and validation of HTPM correlation modules in healthy immune cells

We hypothesized that comparison of HPTM correlation networks for each immune cell type will identify differences in epigenetic regulations between them. To test this hypothesis, we used BR1 (7.39 million cells from 12 subjects) as a “discovery” cohort and BR2, Twins, Stanford, and SA (>20 million cells from 71 subjects) as “validation” cohorts. Pairwise Pearson correlations for all HPTMs pairs for 11 immune cell types (HPCs, pDCs, mDCs, NK, NKT, B, CD4, and CD8 T cells, cMOs, iMOs and ncMOs) were highly conserved between discovery (**Fig. 1A, Fig. 2A**) and 4 independent validation cohorts, irrespective of age and country (**Fig. 1B-E**, **Fig. 2B-E, Methods**). Furthermore, we observed significantly higher variance in pairwise HPTM correlations in HPCs and myeloid cells than lymphoid cells (**Fig. S1A**), suggesting higher epigenetic heterogeneity (**Fig. 1F**, **Fig. 2F, Methods**). These consistent, hitherto unobserved, correlations between HPTMs across multiple independent cohorts provide strong evidence of conserved epigenetic regulatory networks in major immune cell types in healthy individuals.

**Figure 1.**
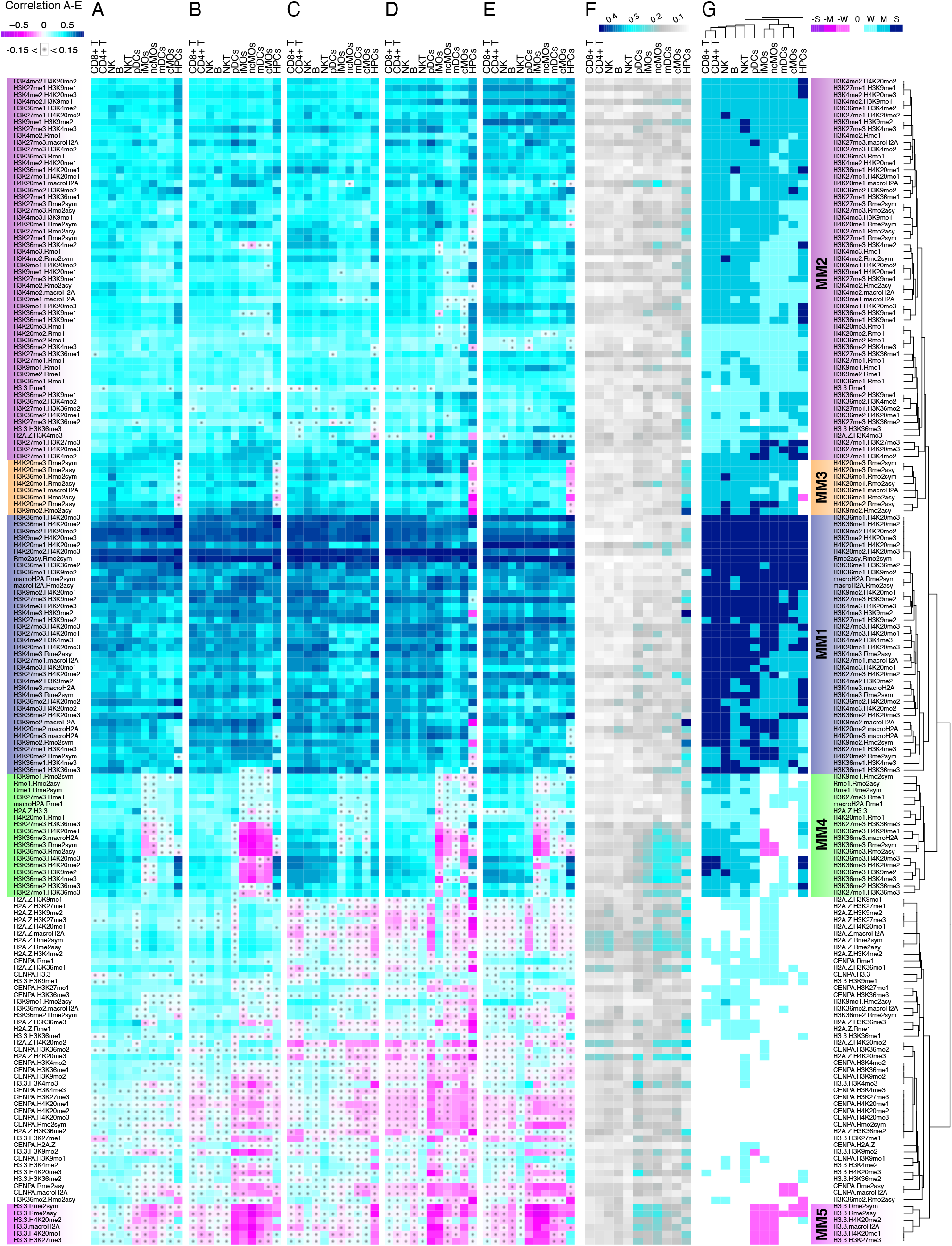
Correlation analysis of the methylation panel shows a high degree of epigenetic conservation across immune cells between independent cohorts. **A-E** Heatmaps show Pearson correlation coefficients between HPTMs for 11 immune cell types in **A** BR1, **B** BR2, **C** Twins, **D** Stanford, and **E** South Africa cohorts. Pairs of HPTMs are rows and immune cell types are columns of the heatmaps. **F** A heatmap of standard variance between Pearson correlation coefficients from BR1, BR2, Twins, Stanford, and South Africa cohorts. Higher variance indicates increased heterogeneity. **G** A heatmap of averaged and stratified Pearson correlation coefficients between HPTMs for 11 immune cell types showing five methylation modules.

**Figure 2.**
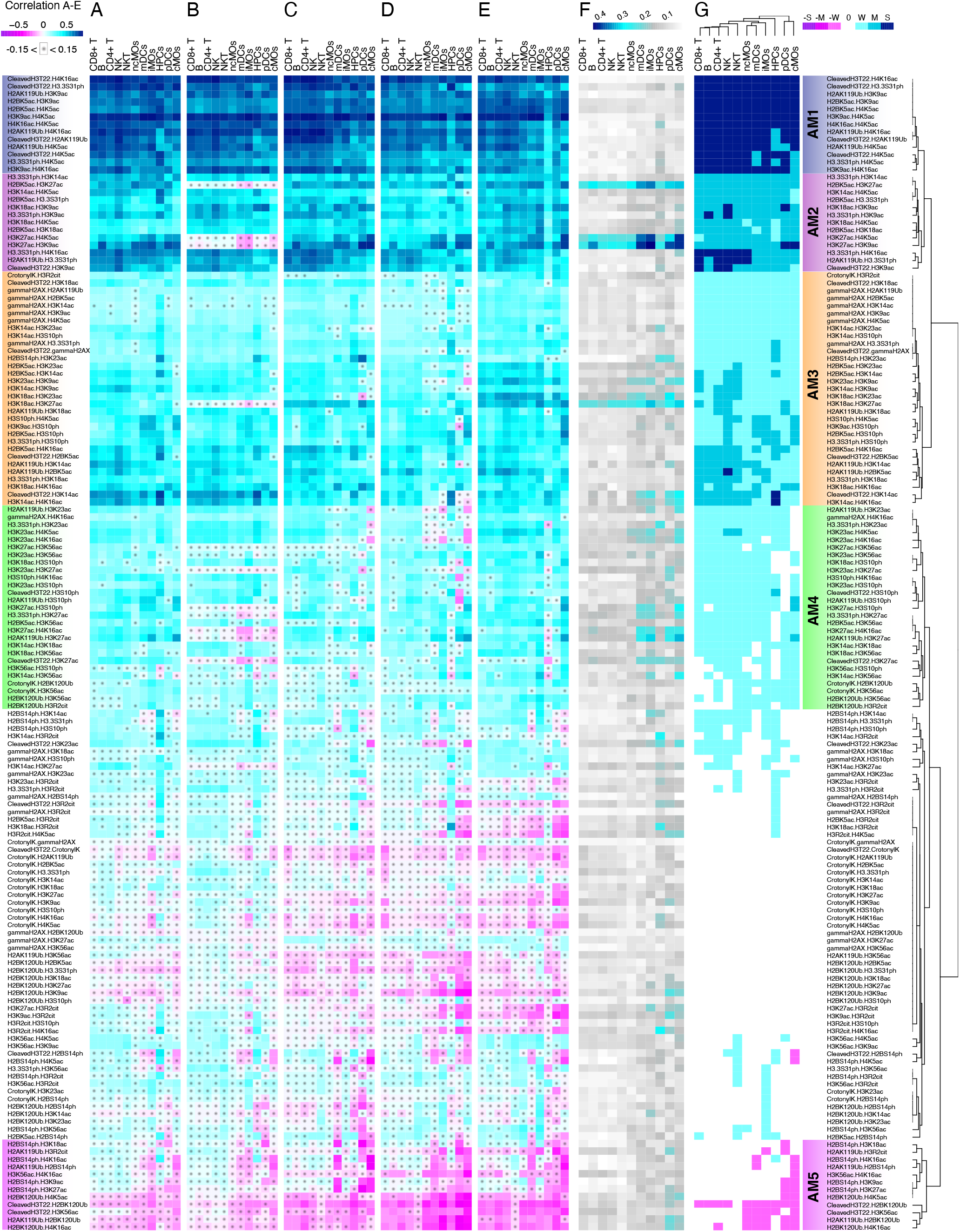
Correlation analysis of the acetylation panel shows a high degree of epigenetic conservation across immune cells between independent cohorts. **A-E** Heatmaps show Pearson correlation coefficients between HPTMs for 11 immune cell types in **A** BR1, **B** BR2, **C** Twins, **D** Stanford, and **E** South Africa cohorts. Pairs of HPTMs are rows and immune cell types are columns of the heatmaps. **F** A heatmap of standard variance between Pearson correlation coefficients from BR1, BR2, Twins, Stanford, and South Africa cohorts. Higher variance indicates increased heterogeneity. **G** A heatmap of averaged and stratified Pearson correlation coefficients between HPTMs for 11 immune cell types showing five acetylation modules.

Given highly reproducible pairwise HPTMs correlations across all independent cohorts, we calculated an average correlation of all HPTM pairs for each immune cell type (**Fig. S1B-C**), stratified the average correlations as strong (|R|≥ 0.6), moderate (0.4 ≤ |R| < 0.6), weak (0.15 ≤ |R| < 0.4), or no correlation (|R| < 0.15), and determined the epigenetic modules in each immune cell type (**Fig. 1G**, **Fig. 2G**, **Methods**). Hierarchical clustering using the average pairwise HPTMs correlations (**Methods**) identified five and four distinct modules in methylation and acetylation panels, respectively, and recapitulated the known hematopoietic differentiation hierarchy in both EpiTOF panels, demonstrating HPCs, myeloid, and lymphoid lineages have distinct correlation networks of HPTMs (**Fig. 1G**, **Fig. 2G**). We assessed the number of connections (N) and betweenness centrality (BC) of the HPTMs within individual modules for each cell type to identify important HPTMs (**Methods**). Betweenness centrality indicates the number of the shortest paths passing through the graph node, i.e., an HPTM, and indicates which node is central to the network.

### Conserved HPTM correlation modules across immune cells

Methylation module 1 (MM1) and 2 (MM2) (**Fig. 3A-B**, **Fig. S2A-B**), and acetylation module 1 (AM1), 2 (AM2), and 3 (AM3) (**Fig. 4A-C**, **Fig. S3A-C**) were present in all immune cell types. MM1 and AM1 were characterized by moderate-to-strong positive correlations, whereas MM2, AM2, and AM3 had weak-to-moderate positive correlations. MM1 comprised 14 HPTMs, of which H3K4me3, H3K9me2, and H4K20me2 were positively correlated with 9 or more HPTMs. Moreover, the highest BC value of H3K4me3 and H3K9me2 indicated their centrality in MM1 (**Fig. 3A**, **Fig. S2A**). These results are in line with previous studies showing that H3K4me3 is enriched within active promoters and regulatory elements^14^, whereas H3K9me2 is modestly correlated with silenced genes and many active genes show H3K9me2 enrichment at their promoters^15^.

**Figure 3.**
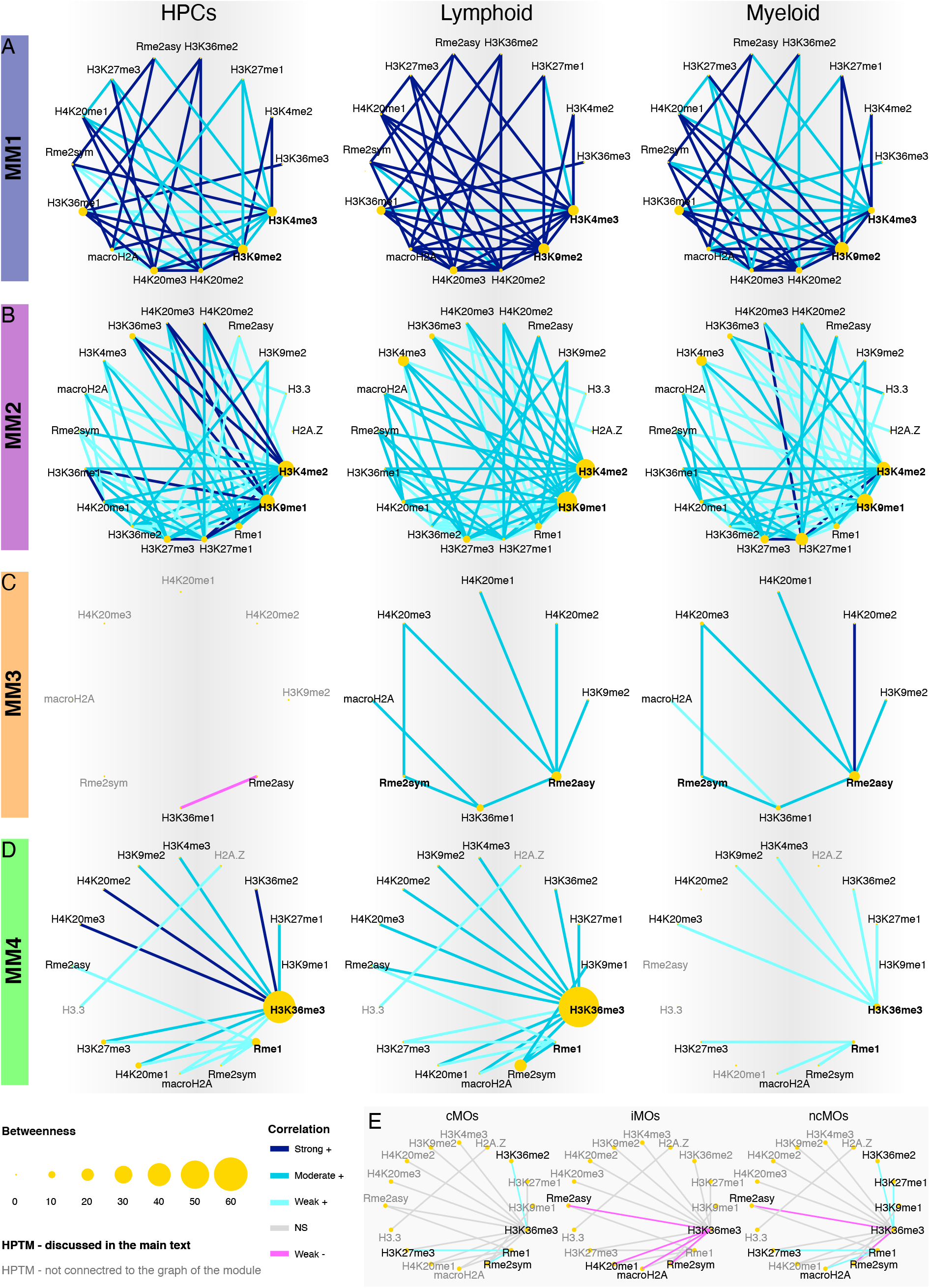
Histone methylation modules in HPCs, average lymphoid and average myeloid cells. **A-B** Graph renderings show **A** methylation module 1 and **B** methylation module 2 that were present in all immune cell types. **C** Methylation module 3 was present in differentiated cells. **D** Methylation module 4 (MM4) differentiated myeloid from lymphoid lineage, mostly because of classical (cMOs), intermediate (iMOs) and non-classical (ncMOs) monocytes. **E** Graph renderings of MM4 in cMOs, iMOs and ncMOs.The first graph in each row **A-D** is the module in HPCs, second is an average representation of lymphoid cells (B, NK, NKT, CD4+, CD8+ T cells) and the third one is an average representation of myeloid cells (cMOs, iMOs, ncMOs, mDCs, pDCs). The color of the edges represents stratified correlation coefficients.

**Figure 4.**
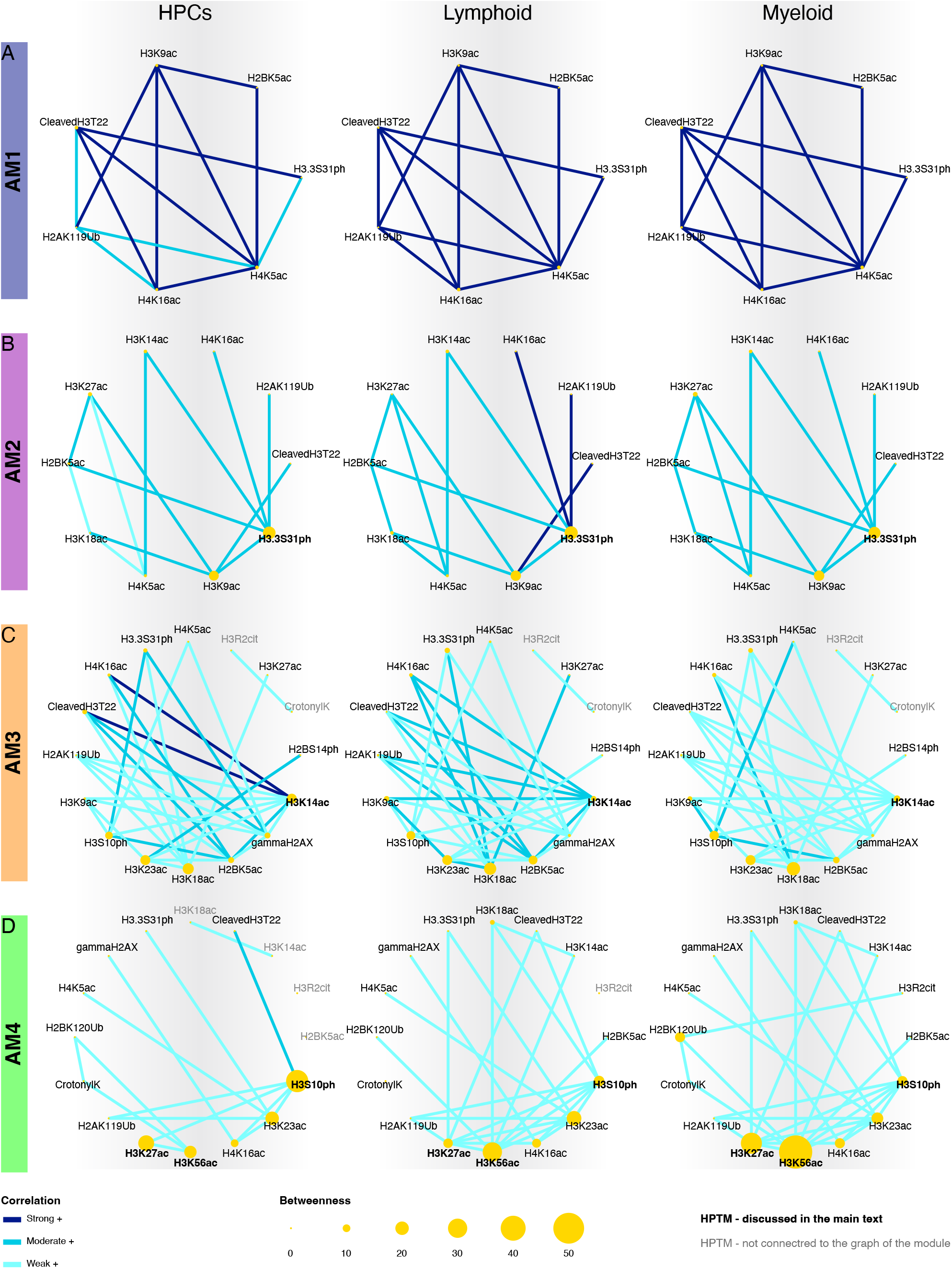
Histone acetylation modules in HPCs, average lymphoid and average myeloid cells. **A-C** Graph renderings show **A** acetylation module 1, **B** acetylation module 2, and **C** acetylation module 3 that were present in all immune cell types. **D** Acetylation module 4 was present in only differentiated cells. The first graph in each row **A-D** is the module in HPCs, second is an average representation of the lymphoid cells (B, NK, NKT, CD4+, CD8+ T cells) and the third one is an average representation of the myeloid cells (cMOs, iMOs, ncMOs, mDCs, pDCs). The color of the edges represents stratified correlation coefficients.

MM2 consisted mostly of weak-to-moderate correlations between 18 HPTMs, of which H3K4me2, H3K9me1, Rme1, H3K27me1, and H3K27me3 were the most connected HPTMs. H3K4me2 and H3K9me1 had the highest betweenness centrality (**Fig. 3B**, **Fig. S2B**), indicating their importance in signal transduction between HPTMs in MM2. Interestingly, MM2 centered around H3K4me2 and H3K9me1, both HPTMs containing one methyl fewer than the central HPTMs in MM1, suggesting methylations of H3K4 and H3K9 may be regulated at the same time.

AM1 was characterized by strong correlations between 7 HPTMs, of which H4K5ac, H4K16ac, H2AK119Ub, and cleaved H3T22 were correlated with each other in all 11 immune cell types (**Fig. 4A**, **Fig. S3A**). AM2 consisted of 10 moderately correlated HPTMs of which H3.3S31ph and H3K9ac had the highest BC values in the AM2 module (**Fig. 4B**, **Fig. S3B**). H4K5ac, H3K9ac and H3.3S31ph were part of both AM1 and AM2. Therefore, these 3 HPTMs were correlated with the highest number of other HPTMs in AM1 and AM2. Interestingly, phosphorylation of H3.3S31 amplifies stimulation-induced transcription, whereas acetylation of H3K9 and H4K5 mediate cellular shift from transcriptional initiation to elongation^16,17^. Thus, our data suggest AM1 and AM2 are strongly associated with transcription regulation.

Although AM3 was present in all immune cell types, lymphoid cells had moderate, whereas myeloid cells had weak correlations between 16 HPTMs (**Fig. 4C**, **Fig. S3C**). H3K14ac had the highest number of connections, whereas H3K18ac had the highest BC value. Both HPTMs are found in transcription start regions of poised and actively transcribed genes^18^, suggesting AM3 is associated with the start of the transcription. Furthermore, since proteolytic cleavage of H3 tails at T22 is involved in gene regulation and physically removes K14 and K18^19^, positive correlations between cleavedH3T22 and H3K14ac or H3K18ac suggest increased acetylations at K14 and K18 occur at untruncated H3 tails, however simultaneously to H3 tail cleavage.

Collectively, presence of the three acetylation (AM1, AM2, and AM3) and two methylation (MM1 and MM2) modules across different immune cells in healthy individuals, irrespective of age and geographic location, suggests their universal importance in immune cells.

### HPTM correlation modules specific to differentiated immune cells

In contrast to the conserved methylation and acetylation modules across all immune cell types, MM3 was present in differentiated immune cells, but not in HPCs (**Fig. 3C**). MM3 included Rme2asy, H3K36me1, Rme2sym, macroH2A, H4K20me1, H4K20me2, H4K20me3, and H3K9me2 with moderate or strong correlations between them in lymphoid and myeloid cells (**Fig. 3C**, **Fig. S2C**). Rme2asy had the highest BC value across all differentiated immune cell types (**Fig. 3C**). The stark difference between HPCs and the differentiated immune cells suggests the presence of MM3 is a hallmark of the differentiated cellular state.

Similarly, only a part of AM4 was present in HPCs (**Fig. 4D**, **Fig. S3D**). This module was characterized by a shift in the highest BC value from H3S10ph in HPCs to H3K56ac in lymphoid and myeloid cells. The shift was associated with an increase in the number of weak correlations for H3S10ph, H3K56ac, and H3K27ac with other HPTMs in lymphoid and myeloid cells. H3K56ac destabilizes the DNA-nucleosome structure and enables unwrapping of the DNA; thus, increasing the binding affinity of the chromatin remodeling proteins that regulate transcription^20^. H3S10ph, among its many functions, regulates transcription by several mechanisms^21^. Together, our results suggest AM4 is associated with establishment of open chromatin and transcription elongation preferentially in differentiated immune cells in blood.

### Lineage-specific HPTM correlation modules

Two methylation modules, MM4 and MM5, were different in myeloid cells compared to HPCs and lymphoid cells. MM4 was present in HPCs and lymphoid cells, but not in all myeloid cells (**Fig. 3D**, **Fig. S2D)**, whereas MM5 was exclusively present in a subset of myeloid cells, centered around H3.3 (**Fig. 1G**). In MM4, H3K36me3 was correlated with the highest number of HPTMs and had the highest BC value in HPCs and lymphoid cells. While H3K36me3 was positively correlated with Rme2sym, macroH2A, H4K20me1, H3K27me3, and Rme2asy in HPCs and lymphoid cells, it had inverse or no correlation with the same HPTMs in cMOs, iMOs and ncMOs (**Fig. 3E**). The negative correlations were driven by the increase in proportions of cMOs, iMOs and ncMOs with higher levels of macroH2A and H3K36me3, rather than the overall increase in abundance of these HPTMs (**Fig. S4**). H3K36me3 marks bodies of recently transcribed genes^22^; thus, our results are in line with transcriptional activity of monocytes during differentiation into ncMOs in blood^23^. In contrast, conserved correlations between H3K36me3 and other HPTMs in MM4 in HPCs and lymphoid cells indicate their low level of transcriptional activity in homeostasis. Collectively, our results identified a putative role of H3K36me3 in regulation of transcription during immune cell differentiation.

Module MM5 consisted of negative correlations between H3.3 and H3K27me3, Rme2sym, Rme2asy, macroH2A, H4K20me1, and H4K20me2 in pDCs, iMOs, and ncMOs (**Fig. 1G**), which is in line with previously described decrease of heterochromatin HPTMs, such as H3K27me3 and macroH2A, with the incorporation of H3.3 into chromatin^24^. H3.3 is enriched in regions of transcribed genes, enhancers, and regulatory elements^25^. Increased proportions of pDCs, iMOs, and ncMOs incorporated H3.3 into chromatin simultaneously increasing H3K36me3 and removing arginine methylations (Rme2sym, Rme2asy), macroH2A (**Fig. S5**), H4K20me1, H4K20me2, and H3K27me3 (data not shown).

MM4 and MM5 demonstrated higher variability in correlations with several HPTMs in myeloid cells. To our knowledge, these modules are the first evidence of dynamic relationships between global arginine methylations (Rme2sym, Rme2asy), H4K20me1 and H4K20me2, and H3K36me3 or H3.3, collectively suggesting during active transcription arginine dimethylations need to be removed.

Overall, our analysis of pairwise HPTM correlation network analysis found highly conserved histone methylations and acetylations modules that fall into three categories: (1) modules conserved across all immune cell types with varying strength of correlations (MM1, MM2, AM1, AM2, and AM3), (2) modules specific to lymphoid (MM4) or myeloid lineage (MM5), and (3) modules specific to differentiated immune cell types but absent or less connected in HPCs (MM3, AM4).

### Peripheral blood hematopoietic progenitors reveal high epigenetic heterogeneity and histone modifications associated with *bona fide* hematopoietic stem cells phenotype

We sought to investigate the extent of epigenetic diversity in an unbiased manner by clustering immune cells using only HPTMs with Phenograph^26^ (**Methods**). When using the methylation panel, we identified 12 cell clusters (**Fig. 5A**), which included (i) 5 methylation clusters (MC1, MC3, MC4, MC5, and MC10) with low abundances of HPTMs, (ii) 1 methylation cluster (MC6) with high abundances of all but 2 HPTMs, (iii) 4 methylation clusters (MC2, MC7, MC8, and MC9) with moderate-to-high abundances of most HPTMs, and (iv) 2 clusters with mixed abundances of HPTMs (MC11 and MC12). As expected from the correlation analysis in which methylation modules MM1, MM2, and MM3 were conserved in all immune cell types (**Fig. 1**), we found each cluster contained every immune cell type (**Fig. 5B**). Despite the conserved methylation modules, we also observed several lineagespecific clusters. For instance, myeloid cells (cMOs, iMOs, ncMOs, mDCs, and pDCs) dominated clusters MC3, MC5, MC7, and MC8 (>75%), whereas lymphoid cells (B, T, NK, and NKT cells) dominated clusters MC1, MC2, MC4, and MC10 (>80%). We also observed clusters dominated by one or two cell types. For instance, cMOs accounted for >50% of MC8, whereas HPCs and pDCs accounted for >75% of MC9 (**Fig. 5B**). MC6 was enriched in HPCs (37.5%), which was marked by higher abundances of H3K9me2, and three methylation states (me1, me2 and me3) of H3K36 and H4K20 (**Fig. 5A**). Notably, higher abundances of these methylation marks in MC6 were solely driven by the HPCs (**Fig. 5C-D**, **Fig. S5**). In fact, average abundance of H3K9me2, H3K36 and H4K20 methylations in 87% of HPCs in MC6 (6.9% of all HPCs) was 2-fold higher that their abundances in the other more differentiated immune cells (**Fig. 5E**, **Fig. S6**). Finally, CD34^dim^CD45^dim^ HPCs separated into two subsets H3K36me1^+^ HPCs (6.9% of total HPCs) and H3K36me1^-^ HPCs (2% of total HPCs), though H3K9me2 was elevated in both subsets (**Fig. 5E**).

**Figure 5.**
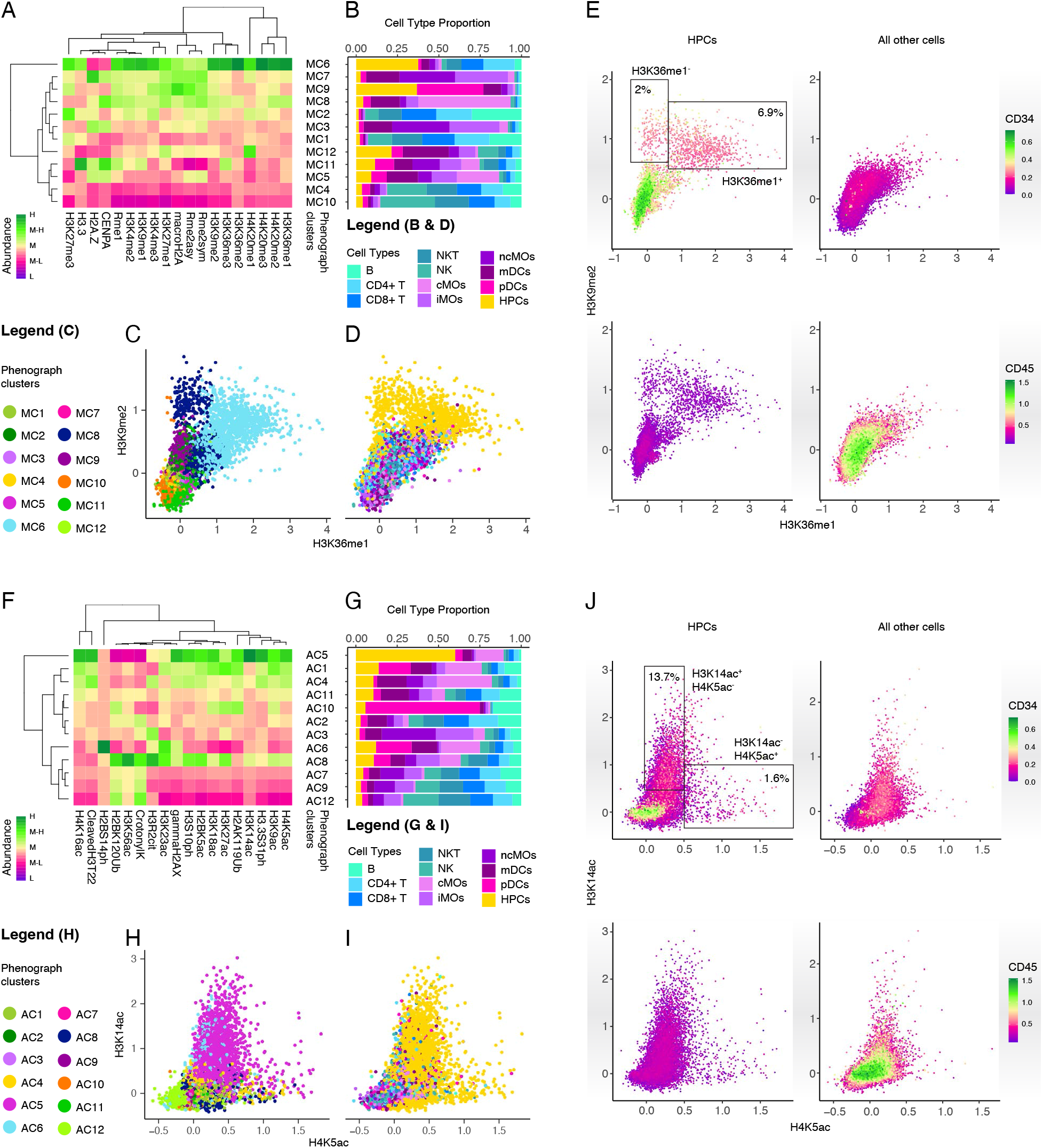
Clustering analysis of methylation and acetylation panels points to increased heterogeneity in hematopoietic cells and reveals distinct patterns of HPTMs. **A** A heatmap showing cell methylation clusters (MC) and average abundances of HPTMs in the methylation panel. Abundance of HPTM is defined in ranges of low (L), moderately low (M-L), moderate (M), moderately high (M-H) and high (H). **B** Proportion of immune cell types in each of the MC clusters. **C-D** Biaxial scatter plots of H3K36me1 and H3K9me2 show all immune cell types labeled for **C** methylation cell clusters (MC) and **D** immune cell types. **C** MC6 cells are high in H3K36me1 and H3K9me2, MC8 cells are low in H3K36me1, while H3K9me2 is high in both clusters. **D** These cells from both clusters (MC6 and MC8) are hematopoietic progenitors (HPCs). **E** Biaxial scatter plots of H3K36me1 and H3K9me2 show hematopoietic progenitors (the first and third quadrants) and all other immune cell types (the second and fourth quadrants) colored by abundance intensity of CD34 (the first and second quadrants) and of CD45 (the third and fourth quadrants). HPCs high in H3K9me2 are dim in CD34 and CD45 resembling phenotypes of *bona fide* hematopoietic stem cells. **F** A heatmap showing cell acetylation clusters (AC) and average abundances of HPTMs in acetylation panel. Abundance of HPTM is defined in ranges of low (L), moderately low (M-L), moderate (M), moderately high (M-H) and high (H). **G** Proportion of immune cell types in each of the AC clusters. **H-I** Biaxial scatter plots of H4K5ac and H3K14ac show all immune cell types labeled for **H** acetylation cell clusters (AC) and **I** immune cell types. **H** AC5 and AC6 cells are high in H3K14ac or H4K5ac. **I** Hematopoietic progenitors (HPCs) have two subpopulations H3K14ac^+^H4K5ac^-^ and H3K14ac^-^H4K5ac^+^. **J** Biaxial scatter plots of H4K5ac and H3K14ac show hematopoietic progenitors (the first and third quadrants) and all other immune cell types (the second and fourth quadrants) colored by abundance intensity of CD34 (the first and second quadrants) and of CD45 (the third and fourth quadrants). H3K14ac^+^H4K5ac^-^ and H3K14ac^-^H4K5ac^+^ HPCs are dim in CD34 and CD45 resembling phenotypes of *bona fide* hematopoietic stem cells.

Similarly, using the acetylation panel, we identified 12 cell clusters of which 8 acetylation clusters (AC2, AC3, AC6, AC7, AC9, AC10, AC11 and AC12) had low to moderate abundances of HPTMs, whereas 4 acetylation clusters (AC1, AC4, AC5, and AC8) had moderate to high abundances of HPTMs (**Fig. 5F**). Overall, we found a mutually exclusive epigenetic pattern shared by all immune cell types. Cell clusters (e.g., AC1, AC4, AC5) with high H4K5ac, H3K9ac, H3.3S31ph, H3K14ac, H2AK119Ub, H3K27ac, H3K18ac, H2BK5ac, H3S10ph, cleaved H3T22, and H4K16ac had low-to-moderately low H3R2cit, lysine crotonylation (crotonylK), H3K56ac, H2BK120Ub, and H2BS14ph. In contrast, cell clusters with higher H3R2cit, H3K56ac, H2BK120Ub, H2BS14ph and crotonylK (e.g., AC7, AC9, AC12) had lower abundances of the other HPTMs (**Fig. 5F**). All acetylation clusters contained every immune cell type, but we also found cell type-specific clusters. For instance, HPCs and pDCs accounted for more than 60% in cluster AC5 and AC10, respectively (**Fig. 5G**). Further, we found higher abundances of HPTMs in AC5 were driven by HPCs and pDCs (**Fig. S7**). Finally, we observed two mutually exclusive subpopulations of peripheral blood CD34^+^ HPCs in AC5 (**Fig. 5H-I**) that were either H3K14ac^+^H4K5ac^-^ (13.7% of total HPCs) or H3K14ac^-^ H4K5ac^+^ HPCs (1.6% of total HPCs). Comparable to the HPCs in methylation panel, the HPCs in AC5 were CD34^dim^CD45^dim^ (**Fig. 5J**), probably representing *bona fide* hematopoietic stem cells.

Across both EpiTOF panels, we identified clusters dominated by HPCs, suggesting higher epigenetic heterogeneity in HPCs, in line with the correlation analyses.

### Lymphoid and myeloid cells follow distinct epigenetic trajectories

Presence of every immune cell type in all clusters defined using HPTMs suggests each immune cell type exists on a continuum. Because histones are post-translationally modified in a coordinated manner in response to stimuli, trajectory inference analysis can identify these coordinated modifications along a continuum. To date, no cellular trajectories using HPTMs have been described.

We used tSpace, a trajectory inference algorithm to define epigenetic trajectories of immune cells (**Methods, Fig. S7A-B**)^27^. When using the methylation panel, tSpace identified a circular trajectory (**Fig. 6A**). We arbitrarily defined the origin of the circular trajectory as the region with the highest proportion of HPCs, which corresponded to the methylation cell cluster MC6, characterized by higher abundances of methylation at H3K9, H3K36, and H4K20 (**Fig. 6B**). Using the origin as a reference, we identified two segments in the circular trajectory that converged on the opposite side. The upper and lower segments were enriched in myeloid and lymphoid cells, respectively (**Fig. 6A, Fig. S8C**). The myeloid segment was characterized by high abundances of all but one histone methylation, H3K27me3, which was higher in the lymphoid segment (**Fig. 6C**). H3K27me3 is a repressive HPTM that enables formation of heterochromatin, therefore suggesting lymphocytes have globally more compact chromatin regions and possibly smaller transcriptional gene repertoire than myeloid cells. However, colocalization of H3K27me3 and H3K4me3 within the promoter regions marks bivalent promoters poised for rapid expression after H3K27me3 removal^28^. Interestingly, H3K4me2 and H3K4me3 were present in high abundance in the lymphoid, but not the myeloid segment (**Fig. 6C**), suggesting the presence of bivalent promoters that allow lymphocytes to circulate blood with minimal gene expression and rapidly increase expression of genes when antigens are detected^29–31^. Collectively, our results are in line with previous studies showing that lymphocytes have higher heterochromatin, small nuclei, and a reduced transcriptional rate compared to myeloid cells^32,33^.

**Figure 6.**
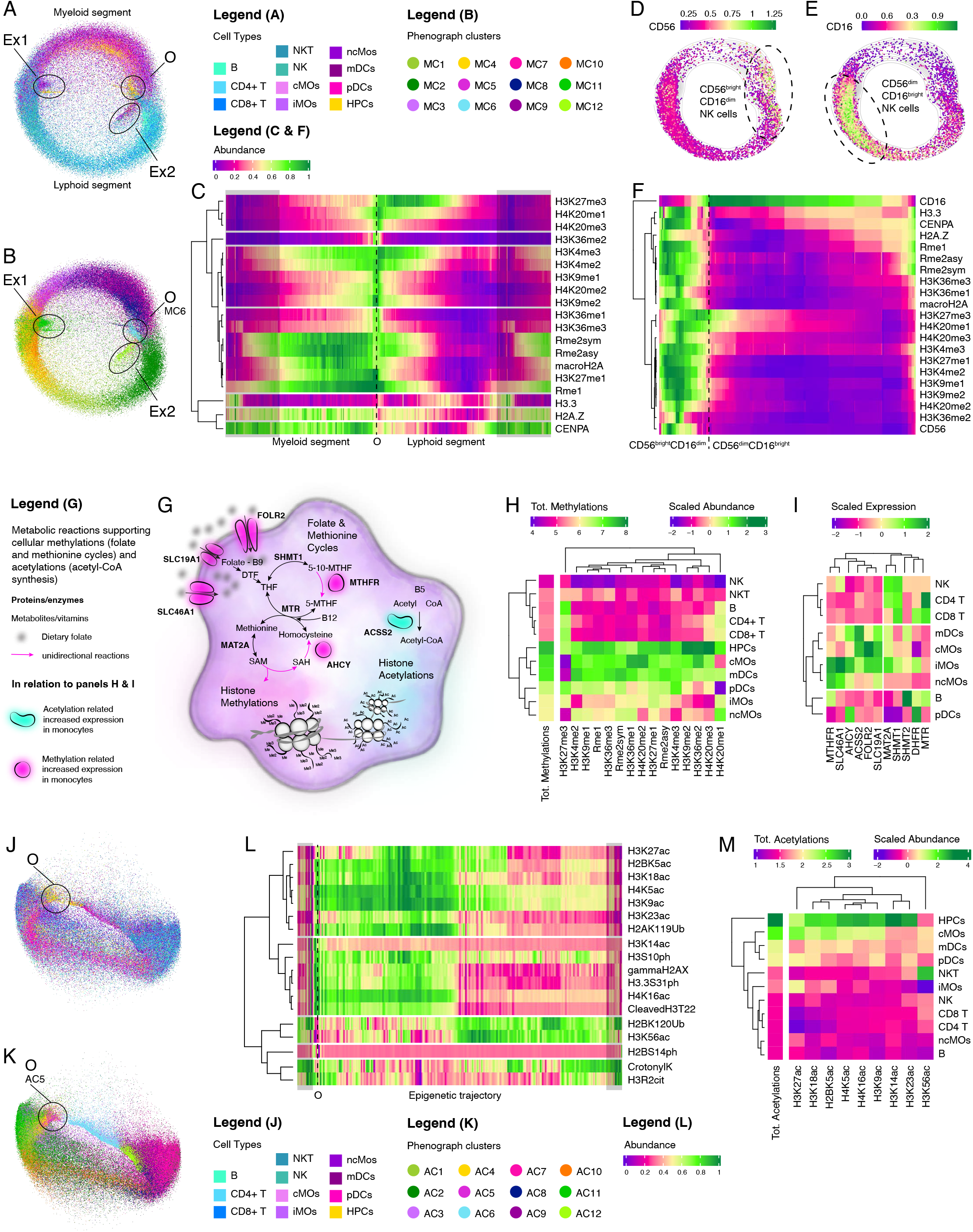
Integration of trajectory inference analysis with Human Blood Atlas transcriptome shows association of HPTM profiles with one-carbon and acetyl-CoA metabolism, cellular lifespan, and immune cell memory. **A-B** Methylation trajectory containing all immune cells colored by **A** immune cell type and **B** cell methylation clusters (MC) shows arbitrarily defined origin of the circular trajectory (O) as the region with the highest proportion of HPCs and MC6 and clusters Ex1 and Ex2. **C** A heatmap of linearized methylation trajectory shows HPTM patterns that define myeloid and lymphoid cell types. Myeloid segment is notably more enriched in histone methylations. Transparent gray rectangles mark the same segments of trajectory, since trajectory is circular. Abundances of HPTMs are scaled 0 to 1. Dashed line marks arbitrarily defined origin (O) of the circular trajectory. **D-E** Methylation trajectory of NK cells colored by abundance of **D** CD56 and **E** CD16 shows **D** CD56^bright^CD16^dim^ and **E** CD56^dim^CD16^bright^ NK cells. **F** A heatmap of linearized methylation trajectory shows NK cells and HPTM patterns associated with CD56^bright^CD16^dim^ and **E** CD56^dim^CD16^bright^ NK cells. CD56^bright^CD16^dim^ NK cells have a high abundance of HPTMs compared to CD56^dim^CD16^bright^ NK cells. Abundances of HPTMs are scaled 0 to 1. Dashed line marks an arbitrary border between CD56^bright^CD16^dim^ and CD56^dim^CD16^bright^ NK cells. **G** An overview of the one-carbon and acetyl-CoA metabolism includes summary of results from panel **I**. Reactions of folate and methionine cycles lead to synthesis of S-adenosylmethionine (SAM) that is a substrate for all cellular methylations. Acetyl-CoA is a substrate for cellular acetylations. One-carbon and acetyl-CoA metabolism depend on vitamins B9 (folate), B12 (cobalamin) and B5 (pantothenic acid). These reactions are the “highway” that connects the environment and global epigenetic changes. S-adenosylhomocysteine (SAH), dihydrofolic acid (DHF), tetrahydrofolic acid (THF), coenzyme A (CoA), folate receptor 2 (*FOLR2*), solute carrier family 19 member 1 (*SLC19A1), SLC46A1*, methylenetetrahydrofolate reductase (*MTHFR*), 5,10-methylenetetrahydrofolate (5-10-MTHF), 5-methyltetrahydrofolate (5-MTHF), 5-methyltetrahydrofolate-homocysteine methyltransferase (*MTR*), adenosylhomocysteinase (*AHCY*), serine hydroxymethyltransferase 1 (*SHMT1*), methionine adenosyltransferase 2A (*MAT2A*), acyl-CoA synthetase short chain family member 2 (*ACSS2*) **H** A heatmap of summed abundances for each histone methylation with sum of all histone methylations in each immune cell type. Individual histone methylations are shown as z-score scaled values. **I** A heatmap of scaled expression values of metabolic genes for each immune cell type from Human Blood Atlas. Monocytes (cMOs, iMOs, ncMOs) and dendritic cells have upregulated critical genes in one-carbon and acetyl-CoA metabolism compared to lymphocytes. **J-K** Acetylation trajectory containing all immune cells colored by **J** immune cell type and **K** cell acetylation clusters (AC) shows arbitrarily defined origin of the circular trajectory (O) as the region with the highest proportion of HPCs and AC5. **L** A heatmap of linearized acetylation trajectory shows HPTM patterns that are shared between all immune cell types. Transparent gray rectangles mark the same segments of trajectory, since trajectory is circular. Abundances of HPTMs are scaled 0 to 1. Dashed line marks arbitrarily defined origin (O) of the circular trajectory. **M** A heatmap of summed abundances for each histone acetylation with sum of all histone acetylations in each immune cell type. Individual histone acetylations are shown as z-score scaled values.

### Reduced histone methylations are associated with memory and longer cell lifespan

We found NK cells were divided into two populations along the methylation trajectory. A population of CD56^bright^CD16^dim^ NK cells localized along the myeloid segment and CD56^dim^CD16^bright^ NK cells localized along the lymphoid segment (**Fig. 6D-E**). CD56^bright^CD16^dim^ NK cells had increased histone methylations in comparison to CD56^dim^CD16^bright^ (**Fig. 6F**). CD56^bright^CD16^dim^ NK cells are cytokine producers with immunoregulatory properties, whereas CD56^dim^CD16^bright^ NK cells exhibit cytotoxic activity and immune memory^34–38^. Separation of functionally distinct NK cell subtypes into two trajectory segments (myeloid and lymphoid) suggests that in addition to distinguishing by lineage, histone methylations may also distinguish metabolically high and low states corresponding to cytokine producing (CD56^bright^CD16^dim^) and dormant memory NK cells (CD56^dim^CD16^bright^), respectively. Indeed, memory CD56^dim^CD16^bright^ NK cells shared methylation trajectory with other memory immune cell types (T and B lymphocytes, **Fig. 6D, Fig. S8C**) and have a longer lifespan than cells without memory (cytokine producing NK cells and myeloid cells)^36,39–42^. This suggests that the methylation trajectories may also be associated with memory and cell lifespan.

We further investigated whether higher histone methylations are associated with cell lifespan. All cellular methylation reactions, including histone modifications, are dependent on S-adenosylmethionine (SAM) as the methyl donor^43^. SAM is replenished in cells through one-carbon metabolism consisting of the folate and methionine cycles (**Fig. 6G**)^44^. Restriction of folate and methionine cycles has been shown to extend cell lifespan^45,46^. Briefly, folate receptor 2 (*FOLR2*), solute carrier family 19 member 1 (*SLC19A1*), and *SLC46A1* transport dietary folate into cells. In folate cycle, methylenetetrahydrofolate reductase (*MTHFR*) converts 5,10-methylenetetrahydrofolate (5-10-MTHF) into 5-methyltetrahydrofolate (5-MTHF), supporting the synthesis of SAM through a reaction catalyzed by 5-methyltetrahydrofolate-homocysteine methyltransferase (*MTR*)^44^. In the methionine cycle, adenosylhomocysteinase (*AHCY*) converts S-adenosyl-L-homocysteine (SAH) into homocysteine, which is a precursor of SAM (**Fig. 6G)**. Both are unidirectional biochemical reactions in the folate and methionine cycles.

Myeloid cells had substantially higher net sum of all methylations than lymphoid cells (**Fig. 6H; Methods**). Therefore, we hypothesized that one-carbon metabolism is increased in myeloid cells. To test this hypothesis, we examined the expression of folate receptors, transporters, and genes encoding for enzymes involved in folate or methionine cycles using The Human Blood Atlas (**Methods**)^47^. Classical, intermediate, and non-classical monocytes had higher expression of receptors (*FOLR2, SLC19A1, SLC46A1*) and two enzymes (*AHCY, MTHFR*) compared to lymphoid cells, suggesting lymphoid cells may have reduced active uptake of dietary folate and reduced synthesis of SAM (**Fig. 6I**)^48^. These results suggest most lymphoid cells have restricted folate and methionine metabolism compared to myeloid cells. Furthermore, lymphoid cells (T, B, and CD56^dim^CD16^bright^ NK cells) with reduced global histone methylations and low expression of crucial enzymes in folate and methionine cycles, live up to several years and exhibit immune memory, whereas myeloid cells, which exhibit upregulated folate and methionine cycles, live up to 14 days^36^. Our findings are in line with studies showing restriction of folate and methionine metabolism extends lifespan^45,46^, but extending them to demonstrate, for the first time, difference in global histone methylation abundances and one-carbon metabolism between myeloid and lymphoid lineages associated with the length of cellular life span in immune cells.

### The highest abundances of H3K36me3 and H4K20me1 are associated with transcriptional activity and differentiation of monocytes

Trajectory analysis also identified two clusters, Ex1 and Ex2 (**Fig. 6A-B**). Cluster Ex1 was enriched in myeloid cells (especially cMOs, iMOs and ncMOs) and characterized by high abundance of H3K36me3 (**Fig. S8D**). Cells increase abundance of H3K36me3 within actively transcribed regions after RNA polymerase II transcribed the region to prevent spurious RNA polymerase II transcription initiation from within the gene bodies^49^. Thus, our findings indicate differentiation of monocytes from cMOs to iMOs to ncMOs is accompanied by the increased transcriptional activity, reflecting ongoing monocytic differentiation in blood^50^.

Cluster Ex2, characterized by a high abundance of H4K20me1, was enriched in mDCs, iMOs, ncMOs, and NK cells (**Fig. S8E**). H4K20me1 is associated with active promoters, initiation of transcriptional activation, and tightly regulated by the cell cycle progression with the highest abundance in mitosis and G1 phase^51^. Therefore, mDCs, iMOs, ncMOs, and NK cells in Ex2 cluster may be recently divided cells that exited the mitosis and are in G0/G1 phase. Separation of Ex1 and Ex2 clusters in trajectory inference analysis indicates cells with the highest abundances of H3K36me3 and H4K20me1 are not abundant in all other HPTMs that were largely enriched along the methylation trajectory.

### Increased histone acetylations in myeloid cells are metabolically supported

A similar trajectory inference analysis of the acetylation panel showed immune cells exist in two mutually exclusive epigenetic states (**Fig. 6J-L, Fig. S8B**). All immune cells harbor either high abundances of most HPTMs with low abundances of H2BK120Ub, H3K56ac, CrotonylK and H3R2cit, or low abundances of most HPTMs with high abundances of H2BK120Ub, H3K56ac, CrotonylK, and H32Rcit.

However, the net sum of all acetylations for each cell type demonstrated a marked increase in multiple histone acetylation marks in HPCs, mDCs, and cMOs compared to B, T, and NK cells (**Fig. 6M**, **Methods**), suggesting myeloid cells exhibit more open chromatin that is permissive to transcription^52^. Moreover, all cellular acetylation reactions depend on acetyl-Coenzyme A (acetyl-CoA) as a substrate, which is synthesized by acetyl-CoA synthetase 2 (*ACSS2*)^53^ *ACSS2* is expressed at a higher level within the myeloid compartment, specifically in cMOs, mDCs, and iMOs (**Fig. 6I**). Further, H3K27ac was higher in cMOs, iMOs, HPCs, and pDCs, which marks enhancers and promoters of actively transcribed genes. Its abundance in monocytic cells indicates activation of gene expression associated with differentiation^50,54^. Strikingly, across all immune cell types H3K56ac was the highest in NKT cells (**Fig. 6M**), which has not been reported before. While H3K56ac has been demonstrated to open chromatin, it is also implicated in DNA damage sensing and response^55^, which stalls DNA replication forks and induce the expression of ligands for the NKG2D receptor found in NK, NKT and CD8+ T cells^56^.

In summary, our analysis revealed cell-specific differences in global histone methylations and acetylations supported through modulations of one-carbon and acetyl-CoA metabolism, respectively. Furthermore, histone methylations separate immune cells according to their ontogeny and are associated with cellular lifespan and memory.

## Discussion

We performed HPTM profiling of almost 28 million PBMCs from 5 independent cohorts of 83 healthy controls, which represented real-world biological heterogeneity, by measuring 37 HPTMs and histone variants across 11 immune cell types. To the best of our knowledge, this is the most comprehensive single cell epigenetic profiling of healthy PBMCs to date. We modelled cross talk between HPTMs using linear and nonlinear analyses to identify (1) HPTM modules conserved across all immune cell types or specific to a cell type, (2) epigenetic heterogeneity at single-cell level, and (3) differences in global histone methylations and acetylations between innate and adaptive immune cells that are associated with metabolism. We found that basal level transcriptional differences in one-carbon metabolism are associated with intrinsic differences in cellular life span and capacity to mount immune memory between innate and adaptive immune cells. Our results establish the reference HPTM landscape of the healthy human immune system and provide the foundation for future studies aimed at identifying perturbed HPTM pathways within immune cells in cancers, vaccines, infections, and autoimmune diseases.

HPTM correlation network analysis identified hub HPTMs, characterized by high betweenness centrality, which suggests that immune cells rely on a handful of hub HPTMs that are highly correlated with the other HPTMs. Irrespective of their different homeostatic physiological functions, each immune cell type shared modules with hub HPTMs involved (1) in activation (H3K4me2/ me3, H3K9me1) and repression (H3K9me2) of gene transcription through chromatin remodeling of transcription start sites (TSSs), or (2) in epigenetic bookmarking (H4K5ac) and regulation of distant enhancers (H3K9ac, H3K14ac and H3.3S31ph)^15,16,57,58^. Therefore, we propose these conserved modules, identified using tens of millions of cells, are strong evidence of conserved coregulation of *trans* and *cis* histone pathways. Importantly, the highest centrality of H3K4me3/H3K9me2 in MM1, H3K4me2/H3K9me1 in MM2, and clustering of these HPTMs in trajectory inference analysis, strongly suggest immune cells regulate transcription through a *cis* bivalent histone methylation signature H3K4me3-K9me3 /2^59–61^. This bivalent signature has not been described before in any immune cell type. Considering the robustness of our results and recent advances in targeting lysine demethylases *KDM7B* and *KDM4A*, which modulate these two HPTMs, with small molecule inhibitors, further studies should focus on understanding their functions^62–64^.

Proteolytic cleavage of H3 histone at threonine 22 (cleaved H3T22) physically removes histone tails including N terminal end up to K27^19^. However, we discovered positive correlations between cleaved H3T22 and H3K9ac, H3S10ph, H3K14ac, H3K18ac, and H3.3S31ph. Of these HPTMs, all but H3.3S31ph cannot occur if the histone tail is physically removed, thus it seems H3K9ac, H3S10ph, H3K14ac, and H3K18ac are introduced in chromatin through independent H3 tails but during proteolytic cleavage. Interestingly, cleaved H3T22 had no or weak correlation with H3K23ac and H3K27ac, suggesting acetylations around the cleavage site (K23, K27) either block proteolysis or H3K23ac and H3K27ac are involved in the establishment of the open chromatin once proteolytic activity decreased. Functional role of acetylation at H3K23, aside from constituting open chromatin, is poorly understood; thus, further investigation is needed especially in conjunction with cleaved H3.

In lineage specific HPTM modules, hub HPTMs were involved in direct regulation of active transcription (e.g., H3K36me3 in MM4 and H3.3 in MM5). Both correlation analysis and trajectory inference found that when cMOs, ncMOs, and iMOs increase H3K36me3 or H3.3, other histone methylation marks are concurrently removed from the histone tails. H3K36me3 prevents random positioning of RNA pol II during transcription and occurrence of faulty transcripts through transcription elongation^49,65,66^. Further, because nucleosomes are assembled in pairs of histone dimers, i.e., H3-H4 and H2A-H2B, it is reasonable to conclude that cells swap modified pairs of canonical H3-H4 with the newly synthetized H3.3-H4 pairs^67^. Such a dramatic chromatin remodeling, mostly in cMOs, ncMOs, and iMOs, has not been reported before, and is likely an indication of their differentiation in blood^50^. Overall, our data support a model in which cells remove other HPTMs that may promote transcription, further suggesting strict compartmentalization between gene regulation and transcription.

We defined HPCs as Lin^-^CD34^+^ PBMCs. However, CD34 is not exclusively expressed in hematopoietic stem cells but is also expressed in more differentiated cells, including CMPs, CLPs, and MEPs^68^. Hence, HPCs in blood are a mixture of different progenitors. Trajectory analysis followed by clustering of PBMCs identified epigenetically distinct subpopulations of HPCs, including H3K36me1^+^ HPCs (6.9% of total HPCs) and H3K36me1^-^ HPCs (2% of total HPCs) using the methylation panel, and H3K14ac^+^H4K5ac^-^ HPCs (13.7% of total HPCs) and H3K14ac^-^H4K5ac^+^ HPCs (1.6% of total HPCs) using the acetylation panel. Importantly, these subpopulations were phenotypically defined as CD45^dim^CD34^dim^ HPCs. Since more pluripotent HPCs are considered CD45^dim^CD34^dim 69,70^, our data suggest high abundances of H3K9me2, H3K36me1, H3K14ac, and H4K5ac in CD45^dim^CD34^dim^ HPCs may be the hallmarks of *bona fide* hematopoietic stem cells. These results further highlight the complementary advantage of EpiTOF compared to ChIP-seq, which cannot identify this heterogeneity in a rare population, and ATAC-seq, which can identify the heterogeneity but not the combination of HPTMs that lead to observed chromatin accessibility.

Trajectory inference analysis, integrated with transcriptome data, strongly associated HPTM profiles with one-carbon metabolism, cellular lifespan, and immune cell memory, several of which have been mechanistically demonstrated in other biological systems^43–46,71–80^. We found myeloid and CD56^bright^CD16^dim^ NK cells have higher histone methylations than T, B or CD56^dim^CD16^bright^ NK cells. Transcriptomics data corroborated myeloid cells with increased histone methylations support high turnover of various histone methylations through SAM synthesis^81^, which is energetically expensive to maintain for a longer time. Difference in lifespan between cytokine producing (CD56^bright^CD16^dim^) and memory (CD56^dim^CD16^bright^) NK cells stems from a necessary expression of recombination activating 1 (*RAG1*) and *RAG2* in longer-lived memory, but not cytokine producing NK cells^36,37^. *RAG1* and *RAG2* are also essential for the differentiation of B and T cells, both of which have memory and longer lifespan than other immune cells^34–36,44^. Interestingly, studies in fish found *Rag1*-immunodeficiency induces premature aging and shortens life span, supporting that *RAG1/RAG2* have a role in extension of the life span^82^. However, existing evidence is insufficient to decipher whether the state of low histone methylations is in response to or a driving factor behind the extended life span, warranting future studies addressing mechanisms and causations.

Our study has a few limitations. First, our analysis inferred several novel *cis* and *trans* associations between HPTMs and confirmed several previously described associations and causal relations between HPTMs. These novel associations should be further investigated in mechanistic studies. Reproducibility of these novel associations and cellular heterogeneity across five independent cohorts provide robust evidence in support of further mechanistic studies. For instance, we recently reported epigenetic mechanisms of monocyte differentiation into macrophages, which demonstrated relationships between several HPTMs^83^. Second, EpiTOF lacks locus-specific information, unlike ChIP-seq and ATAC-seq. However, integration of EpiTOF, scATAC-seq and CITE-seq/ scRNA-seq would enhance our understanding of epigenetic processes by identifying a combination of HPTMs that is associated with chromatin accessibility or transcriptome profile in a given cell type during different immunological states. Hence, EpiTOF is an important tool to start mapping at single cell resolution multilayered information coming from chromatin remodeling, transcriptomics, and proteomics.

Overall, our study has long-term implications for immunology, developmental biology, and epigenetics. Our data demonstrated that HPTMs are regulated globally within a cell in a modular fashion, which can be further studied using epigenetic trajectory inference to investigate order of histone modifications in a cell type-specific manner at s single-cell resolution. Most importantly, our study provides a high-resolution reference landscape to start decoding the histone language of the healthy human immune system.

## Contributions

PK conceived the study. PK and PJU obtained funding. DD, LK, AG, and MDo performed computational analyses, and interpreted data. PJU supervised EpiTOF profiling. DD and PK wrote the manuscript with contributions from all coauthors. AK, PC, SC, and MDv performed EpiTOF profiling. TJS enrolled adolescents in South Africa. AH enrolled adults in the Stanford cohort. PK and PJU supervised the study. All authors approved the manuscript.

## Disclosures

PK is funded in part by the Bill and Melinda Gates Foundation (OPP1113682); the National Institute of Allergy and Infectious Diseases (NIAID) grants 1U19AI109662, U19AI057229, and 5R01AI125197; Department of Defense contracts W81XWH-18-1-0253 and W81XWH1910235; and the Ralph & Marian Falk Medical Research Trust. PJU is supported in part by the Donald E. and Delia B. Baxter Foundation, Elizabeth F. Adler, the Henry Gustav Floren Trust, the Bill & Melinda Gates Foundation (OPP1113682), and the NIH grants U19 AI110491 (Autoimmunity Center of Excellence), R01 AI125197.

## Notes

### Competing Interest Statement

The authors have declared no competing interest.

