## Supplementary Materials & Methods for "Crosstalk of Histone Modifications in the Healthy Human Immune System"

**Dermadi *et al.* 2022**

### Supplemental Documentation

### Materials and methods

#### Human Subjects

Written informed consent was obtained from all adult donors in the cohorts and the study was conducted in accordance with the guidance of Stanford Research Compliance Office for Human Subject Research. The protocols were approved by Stanford Institutional Review Board, Autoimmunity Center of Excellence at Stanford (BR1, BR2 and Twins cohorts: IRB-30494; Stanford cohort: IRB- 28427; South Africa cohort: IRB- 41197). Adolescent donors in the South Africa cohort provided written informed assent, while their parents or legal guardians provided written informed consent. The protocol was also approved by the Human Research Ethics Committee of the University of Cape Town.

Detailed information for each cohort can be found in **Table S1**. BR1, BR2 and Twin cohorts were previously published[1]. All participants in biological replicate 1 (BR1), BR2 and Twin cohorts were CMV-seronegative. The BR1 cohort consisted of 12 healthy donors, 6 under 25 and 6 over 65 years (mean age 46), evenly distributed for sex. The BR2 cohort shared number of donors, CMV status and demographics with BR1 (mean age 44). The Twins cohort was balanced in sex and consisted of 8 fraternal and 12 identical twins with an average age of 44.5 years. The Stanford cohort contained 4 females and 5 males with an average age of 40.5 years. The South Africa cohort contained 4 females and 6 males with an average age of 14.8 years.

#### PBMC Isolation

Mononuclear cells were purified from buffy coat by density gradient centrifugation using Ficoll-Paque Plus (GE Healthcare) in SepMate tubes (STEMCELL Technology). Crude PBMCs were treated with RBC lysis buffer (BioLegend) for 5 minutes to remove residual red blood cells followed by 3 PBS washes. Aliquots of 20 million PBMCs were resuspended in human AB serum (Valley Biomedical) containing 5% DMSO (Sigma) for cryopreservation.

#### Lanthanide Labeling of Antibodies for EpiTOF and Panel Design

EpiTOF was developed using lanthanide-based antibody tagging chemistry. We previously characterized and validated 40 commercial highly specific antibodies that recognize different HPTMs, which were distributed in two EpiTOF panels. Briefly, HPTM antibodies were conjugated using MAXPAR antibody labeling kit (Fluidigm), following the manufacturer's protocol. TCEP (Thermo Fisher) was added in 100-fold molar excess to generate sulfhydryl groups for maleimide-mediated conjugation of metal-chelating polymers. Conjugated antibodies were diluted in antibody stabilizing solution (Boca Scientific) containing 0.05% sodium azide (Sigma) for storage.

Two panels share 10 antibodies against phenotypic cell surface markers for immune cells of human peripheral blood and 2 antibodies targeting total histone proteins H3 and H4. One panel detects selected histone methylations, and another selected histone acetylations, phosphorylations, and ubiquitinations (**Table S2.**). Additionally, each panel has a capacity to accommodate up to 20 unique samples e.g., patients or human donors, by utilizing a barcode combination of 6 palladium channels (Fluidigm kit).

#### EpiTOF (Sample Processing, Staining, Barcoding and Data Collection)

Cryopreserved PBMCs were thawed and incubated in RPMI 1640 media (Thermo Fisher) containing 10% FBS (ATCC) at 37° C for 1 hour prior to processing. Cisplatin (ENZO Life Sciences) was added to 10 mM final concentration for viability staining for 5 minutes before quenching with CyTOF Buffer (PBS (Thermo Fisher) with 1% BSA (Sigma), 2mM EDTA (Fisher), 0.05% sodium azide). Cells were centrifuged at 400 g for 8 minutes and stained with lanthanide-labeled antibodies against immunophenotypic markers in CyTOF buffer containing Fc receptor blocker (BioLegend) for 30 minutes at room temperature (RT). Following extracellular marker staining, cells were washed 3 times with CyTOF buffer and fixed in 1.6% PFA (Electron Microscopy Sciences) at $1x10^6$ cells/ml for 15 minutes at RT. Cells were centrifuged at 600 g for 5 minutes post-fixation and permeabilized with 1mL ice-cold methanol (Fisher Scientific) for 20 minutes at 4° C. 4 mL of CyTOF buffer was added to stop permeabilization followed by 2 PBS washes. Sample were barcoded using Fluidigm kit (Cell-IDTM 20-Plex) and following the manufacturer's protocol (Fluidigm). Individual samples were then combined and stained with intracellular antibodies in CyTOF buffer containing Fc receptor blocker (BioLegend) overnight at 4° C. The following day, cells were washed twice in CyTOF buffer and stained with 250 nM 191/193 Ir DNA intercalator (Fluidigm) in PBS with 1.6% PFA for 30 minutes at RT. Cells were washed twice with CyTOF buffer and once with double-deionized water (ddH2O) (Thermo Fisher) followed by filtering through 35 mm strainer to remove aggregates. Cells were resuspended in ddH2O

**Table S1. List of cohorts profiled using EpiTOF**

| Cohort | Donors | Sex | Mean Age (year range) | Number of cells | | Total cells |
|---|---|---|---|---|---|---|
| | | | | Methylation panel | Acetylation panel | |
| BR1 | 12 | 6 males, 6 females | 46 (17-80) | 3,931,571 | 3,466,454 | 7,398,025 |
| BR2 | 12 | 6 males, 6 females | 44 (17-77) | 2,360,243 | 2,360,255 | 4,720,498 |
| Twins | 40 | 10 males, 10 females | 44.5 (16-71) | 4,177,839 | 4,959,450 | 9,137,289 |
| Stanford Cohort | 9 | 5 males, 4 females | 40.5 (27-65) | 2,072,841 | 1,993,232 | 4,066,073 |
| South Africa Cohort | 10 | 6 males, 4 females | 14.8 (13-18) | 1,259,959 | 1,259,959 | 2,519,918 |
| 5 cohorts | 83 | 33 males, 30 females | 37.9 (13-80) | 13,802,453 | 14,039,350 | 27,841,803 |

containing four element calibration beads (Fluidigm) and analyzed on CyTOF2 (Fluidigm) in Stanford Shared FACS Facility. Raw data were concatenated and normalized using calibration beads following the manufacturer's protocol for downstream processing.

#### Immune Cell Population Definitions (Gating Strategies) and Data Pre-Processing

We de-barcoded cell events from individual samples and segregated 11 immune cell populations following a previously published gating strategy in FlowJo[1]. Immune cell types included were hematopoietic progenitor cells (HPCs), plasmacytoid dendritic cells (pDCs), myeloid dendritic cells (DCs), natural killer (NK) cells, NKT, B, CD4, and CD8 T cells, classical monocytes (cMOs), intermediate monocytes (iMOs) and non-classical monocytes (ncMOs). Single-cell data for immune cell types from individual subjects were exported from FlowJo for downstream computational analyses.

#### Data processing and normalization

The data were normalized and processed in a three-step process. First, the raw EpiTOF data was transformed using the formula:

$$HPTM_{transformed} = \frac{\sqrt{HPTM_{raw}}}{50}$$

We then applied a multivariate linear regression to each post-translational modification independently, using all cells in a single sample. The formula for this can be represented as such:

$$HPTM_{i,j} = \beta_0 + \beta_1 H3_i + \beta_2 H4_i$$

Where H3 and H4 are the transformed abundances of H3 and H4 in cell *i*, and $HPTM_{i,j}$ is the transformed value for a given HPTM *j* in cell *i* across all cell types in a single sample. This was done to remove the effect of H3 and H4 on the HPTM. The normalized value for an HPTH can be represented as a residual: the measured $HPTM_{i,j}$ value vs. expected value based on abundance of H3 and H4.

$$HPTM_{i,j\,final} = HPTM_{i,j} - \widehat{HPTM}_{i,j}$$

Finally, instead of using ordinary least squares to find $\beta_0$, $\beta_1$, and $\beta_2$, we used a weighted-least squares regression. We assigned a weight to each cell based on its cellular identity. The weight is calculated as the inverse count of the cell type, $1/n_{celltype}$. If a cell type is not abundant and its count is less than the 3rd quantile of counts from all cell types in a sample, we represent $n_{celltype}$ as the 3rd quantile of cell type counts. This is to allow the regression to approximate an average blood cell, taking into account rarer cell types (e.g., some of myeloid cell types or HPCs), instead of just normalizing to abundant cell types (e.g., CD4 or CD8 T cells).

#### Correlation analysis

Correlation analysis was applied separately to methylation and acetylation panels of each cohort to determine the strength of association between two HPTMs. Using all cells in a respective cohort, we calculated Pearson correlation coefficient for all pairs of HPTMs in an EpiTOF panel for each cell type separately. These correlations between HPTM pairs were visualized as heatmaps using NMF R package (**Fig. 1A-E, Fig. 2A-E**). Standard deviation of HPTM pair correlation coefficients was used to assess the variability of HPTM pairs between the cohorts (**Fig. 1F, Fig. 2F**).

To determine HPTM correlation modules we first calculated average values of HPTM pair correlation coefficients using data from all 5 cohorts. These average correlation coefficients were then stratified into strong ($|R| \geq 0.6$), moderate ($0.4 \leq |R| < 0.6$), weak ($0.15 \leq |R| < 0.4$), or no correlation ($|R| < 0.15$). Hierarchical clustering was used to determine the epigenetic modules. To determine lineage-specific average representation of each module (**Fig. 3 & 4**) we calculated mean of stratified HPTM pair correlation coefficients using CD4, CD8 T cells, NKT and NK cells for lymphoid, and cMOs, iMOs, ncMOs, DCs and pDCs for myeloid lineage. Epigenetic modules were visualized as graph networks using igraph package in R. Every network contains nodes and edges. In our case nodes were HPTMs, and edges were correlation coefficients. In network analysis betweenness centrality is a measure that quantitates how much information passes through a specific node in the network. In other words, high betweenness centrality for specific HPTM indicated its importance in each module. We calculated betweenness centrality using a function in igraph package.

#### Clustering and trajectory inference

Prior to clustering we combined data from all cohorts and randomly downsampled number of cells of every cell type to match the cell number of the smallest cell type, in our case hematopoietic progenitor cells (N = 24,131 in acetylation and N = 11,147 in methylation panel). This balanced yet random downsampling permits exploring of epigenetic changes within immune system that will not be outweighed by most dominant immune cell types. In methylation panel downsampling yielded 122,617 cells across all immune cell types, and in acetylation panel 265,441 cells. These downsampled data were used for further clustering and trajectory inference analysis.

To explore epigenetic heterogeneity within immune system we clustered all immune cell types together using only normalized abundances of HPTMs and Rphenograph package – an R implementation of Phenograph clustering algorithm (**Fig. 5A**)[2]. Phenograph clustering was performed using hyperparameter k of 750 or 1000, for acetylation or methylation panels, respectively. Average values of HPTMs for each cluster were scaled and visualized in heatmaps (NMF R package). We further quantified proportions of different immune cell types for each cluster

Epigenetic trajectories were inferred using all normalized HPTMs in a respective panel and tSpace[3] algorithm with hyperparameters K = 15, T = 500. Methylation and acetylation trajectories were both circular and we isolated them by fitting a principal line using package princurv in R with hyperparameters thresh = 0.053 and smoother = "periodic_lowess" (**Fig. S8A-B**). Abundance of every HPTM was smoothed along the fitted principal line using smooth. spline function in R package stats and scaled between 0 and 1. For visualization purposes we copied one end of the circular trajectory to the other end and labelled copied segment in the figures with a transparent shade of gray (**Fig. 6C & L**). To examine which parts of epigenetic trajectories are occupied by individual immune cell type we visualized epigenetic trajectory for every immune cell type separately (**Fig. S8C**).

However, in methylation panel, prior to trajectory isolation, we removed cells from two clusters that were outside the circular trajectory Ex1 and Ex2. For Ex1 we selected cells that had tPC1 and tPC2 coordinates in the range $-0.003 < tPC1 \leq 0.002$ & $-0.00021 \leq tPC2 \leq 0.0012$, and for Ex2 cells that had coordinates in the range $0.0015 < tPC1 \leq 0.003$ and $-0.0025 \leq tPC2 \leq -0.0008$. These clusters were further analyzed by calculating average abundance of

HPTMs and phenotypic surface marks for each immune cell type in the cluster and visualized as heatmaps (**Fig. S8D & E**). We also calculated cell proportions in each of the clusters.

#### Total histone methylations and acetylations in immune cell types

Trajectory inference pointed myeloid cells have higher methylations than lymphoid cells. To investigate which immune cell types on average have more of any histone methylation or acetylation, we summed abundances of individual histone methylations or acetylations within each immune cell type and visualized scaled values in a heatmap (**Fig. 6H & M**). Then we calculated total methylations or total acetylations by summing together individual sums of histone methylations or histone acetylations for each immune cell type.

#### One carbon metabolism gene expression in immune cell types

Since we detected marked difference in total methylations and acetylations within immune cell types, we investigated gene expressions of receptors and enzymes involved in one carbon and acetyl-CoA metabolism. We mined Human Blood Atlas[4] for expression of *FOLR2*, *SLC19A1*, *SLC46A1*, *AHCY*, *MTHFR*, *DHFR*, *MTR*, *SHMT1*, *SHMT2*, *MAT2A*, *ACSS2* in blood immune cell types. Gene expressions were scaled and visualized in a heatmap (**Fig. 6I**).

### Table S2. EpiTOFPanel Designs

| Acetylation Panel | | | | |
|---|---|---|---|---|
| **Metal** | **Marker** | **Manufacturer** | **Type** | **Clone** |
| 89Y | CD45 | Fluidigm | Mouse IgG1 | HI30 |
| 141Pr | H3 | CST | Rabbit mAb | D1H2 |
| 142Nd | γ-H2AX | CST | Rabbit mAb | 20E3 |
| 143Nd | H2BK5ac | CST | Rabbit mAb | D5H1S |
| 144Nd | H3S10ph | Active Motif | Mouse IgG1 | MABI 0312 |
| 145Nd | CD4 | BioLegend | Mouse IgG1 | RPA-T4 |
| 146Nd | CD8 | BioLegend | Mouse IgG1 | SK1 |
| 147Sm | H4K5ac | Active Motif | Mouse IgG1 | MABI 0405 |
| 148Nd | CD34 | BD | Mouse IgG1 | 8G12 |
| 149Sm | Cleaved H3 (Thr22) | CST | Rabbit mAb | D7J2K |
| 150Nd | H3.3S31ph | Active Motif | Mouse IgG2b | 1A8G10 |
| 151Eu | H3K23ac | RevMab | Rabbit mAb | RM169 |
| 152Sm | H3K9ac | Active Motif | Mouse IgG2a | 2G1F9 |
| 153Eu | H2BS14ph | CST | Rabbit mAb | D67H2 |
| 154Sm | H2AK119ub | CST | Rabbit mAb | D27C4 |
| 155Gd | CD11c | BioLegend | Mouse IgG1 | Bu15 |
| 156Gd | H3K18ac | RevMAb | Rabbit mAb | RM166 |
| 158Gd | H3K56ac | Active Motif | Mouse IgG1 | 12.1 |
| 159Tb | CD14 | BioLegend | Mouse IgG2a | M5E2 |
| 160Gd | PADI4 | OriGene | IgG2a | OTI4H5 |
| 161Dy | H2BK120ub | CST | Rabbit mAb | D11 |
| 162Dy | Crotonyl-Lys | PTM Biolabs | Mouse IgG | 4D5 |
| 163Dy | H3R2cit | Abcam | Rabbit mAb | EPR17703 |
| 164Dy | H3K14ac | CST | Rabbit mAb | D4B9 |
| 165Ho | H3R2/R8/R17cit | Abcam | Rabbit pAb | cat# ab5103 |
| 166Er | CD33 | BioLegend | Mouse IgG1 | WM53 |
| 167Er | CD16 | BioLegend | Mouse IgG1 | B73.1 |
| 168Er | H4K16ac | CST | Rabbit mAb | E2B8W |
| 169Tm | CD123 | BD | Mouse IgG1 | 9F5 |
| 170Er | CD3 | BioLegend | Mouse IgG1 | UCHT1 |
| 171Yb | CD38 | Biolegend | Mouse IgG1 | HIT2 |
| 172Yb | CD56 | BD | Mouse IgG2b | NCAM16.2 |
| 173Yb | H4 | Abcam | Mouse IgG1 | ab31830 |
| 174Yb | H3K27ac | Active Motif | Mouse IgG1 | MABI 0309 |
| 175Lu | CD19 | BioLegend | Mouse IgG1 | HIB19 |
| 176Yb | HLA-DR | BioLegend | Mouse IgG2a | L243 |

| Methylation Panel | | | | |
|---|---|---|---|---|
| **Metal** | **Marker** | **Manufacturer** | **Type** | **Clone** |
| 89Y | CD45 | Fluidigm | Mouse IgG1 | HI30 |
| 141Pr | H3 | CST | Rabbit mAb | D1H2 |
| 142Nd | Arg-me1 | Abcam | Mouse IgG1 | 5D1 |
| 143Nd | Arg-me2 (sym) | CST | Rabbit mAb | 13222 |
| 144Nd | H3K4me2 | Active Motif | Mouse IgG1 | MABI 0303 |
| 145Nd | CD4 | BioLegend | Mouse IgG1 | RPA-T4 |
| 146Nd | CD8 | BioLegend | Mouse IgG1 | SK1 |
| 147Sm | H3K9me2 | Biolegend | Mouse IgG1 | 5E5-G5 |
| 148Nd | CD34 | BD | Mouse IgG1 | 8G12 |
| 149Sm | H3K9me1 | Biolegend | Mouse IgG1 | 7E7.H12 |
| 150Nd | H3K36me3 | RevMab | Rabbit mAb | RM155 |
| 151Eu | H3K27me1 | Active Motif | Mouse IgG2a | MABI 0321 |
| 152Sm | Arg-me2 (asy) | CST | Rabbit mAb | 13522 |
| 153Eu | H3K36me2 | Active Motif | Mouse IgG1 | MABI 0332 |
| 154Sm | H3K27me2 | Active Motif | Mouse IgG2a | MABI 0324 |
| 155Gd | CD11c | BioLegend | Mouse IgG1 | Bu15 |
| 156Gd | H4K20me2 | Active Motif | Mouse IgG1 | MABI 0422 |
| 158Gd | H3.3 | Abcam | Rabbit mAb | EPR17899 |
| 159Tb | CD14 | BioLegend | Mouse IgG2a | M5E2 |
| 160Gd | H4K20me3 | BioLegend | Mouse IgG1 | 6F8-D9 |
| 161Dy | Macro-H2A | Millipore | Mouse IgG2b | 14G7 |
| 162Dy | H3K4me3 | Life | Rabbit IgG | G.532.8 |
| 163Dy | H2A.Z | Abcam | Rabbit mAb | [EPR6171(2)(B)] |
| 164Dy | H3K36me1 | Abcam | Rabbit mAb | EPR16993 |
| 165Ho | H3K27me3 | Active Motif | Mouse IgG1 | MABI 0323 |
| 166Er | CD33 | BioLegend | Mouse IgG1 | WM53 |
| 167Er | CD16 | BioLegend | Mouse IgG1 | B73.1 |
| 168Er | H4K20me1 | Active Motif | Mouse IgG | 5E10-D8 |
| 169Tm | CD123 | BD | Mouse IgG1 | 9F5 |
| 170Er | CD3 | BioLegend | Mouse IgG1 | UCHT1 |
| 171Yb | CD38 | Biolegend | Mouse IgG1 | HIT2 |
| 172Yb | CD56 | BD | Mouse IgG2b | NCAM16.2 |
| 173Yb | H4 | Abcam | Mouse IgG1 | ab31830 |
| 174Yb | CENP-A | MBL | Mouse IgG1 | 3-19 |
| 175Lu | CD19 | BioLegend | Mouse IgG1 | HIB19 |
| 176Yb | HLA-DR | BioLegend | Mouse IgG2a | L243 |

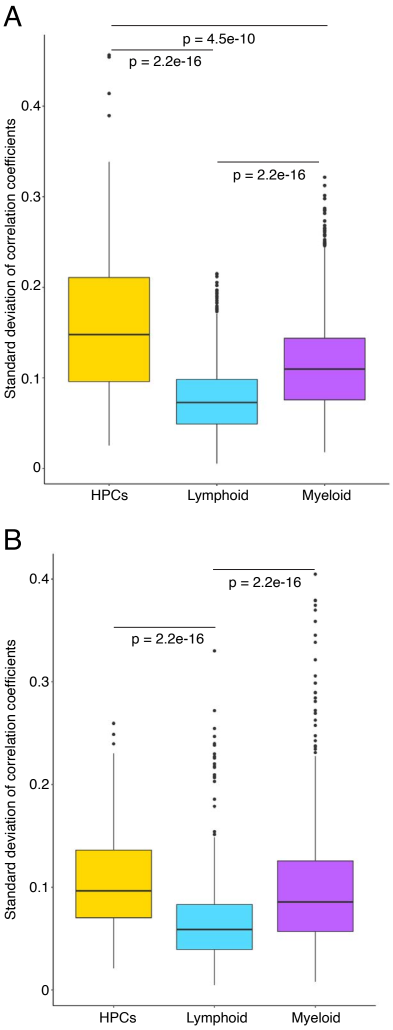


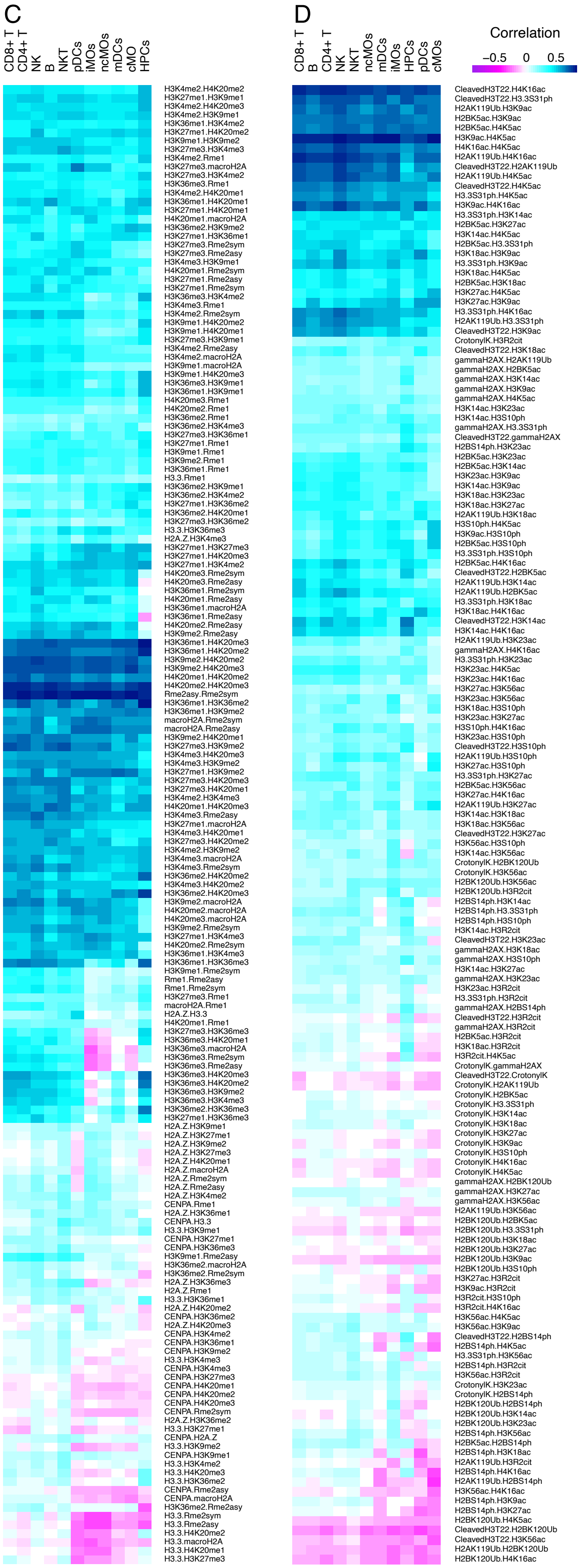


**Fig S1.** (Refers to Figure 1 and 2) Correlation analysis between HPTMs. **A-B** Boxplot of standard deviations of correlation coefficients between HPTM pairs in hematopoietic progenitor cells, cells of lymphoid and myeloid lineages in **A** methylation and **B** acetylation panels. Only significant comparisons are labeled. **C-D** Heatmaps of averaged Pearson correlation coefficients between HPTMs for 11 immune cell types prior to stratification of coefficients in **C** methylation and **D** acetylation panels.

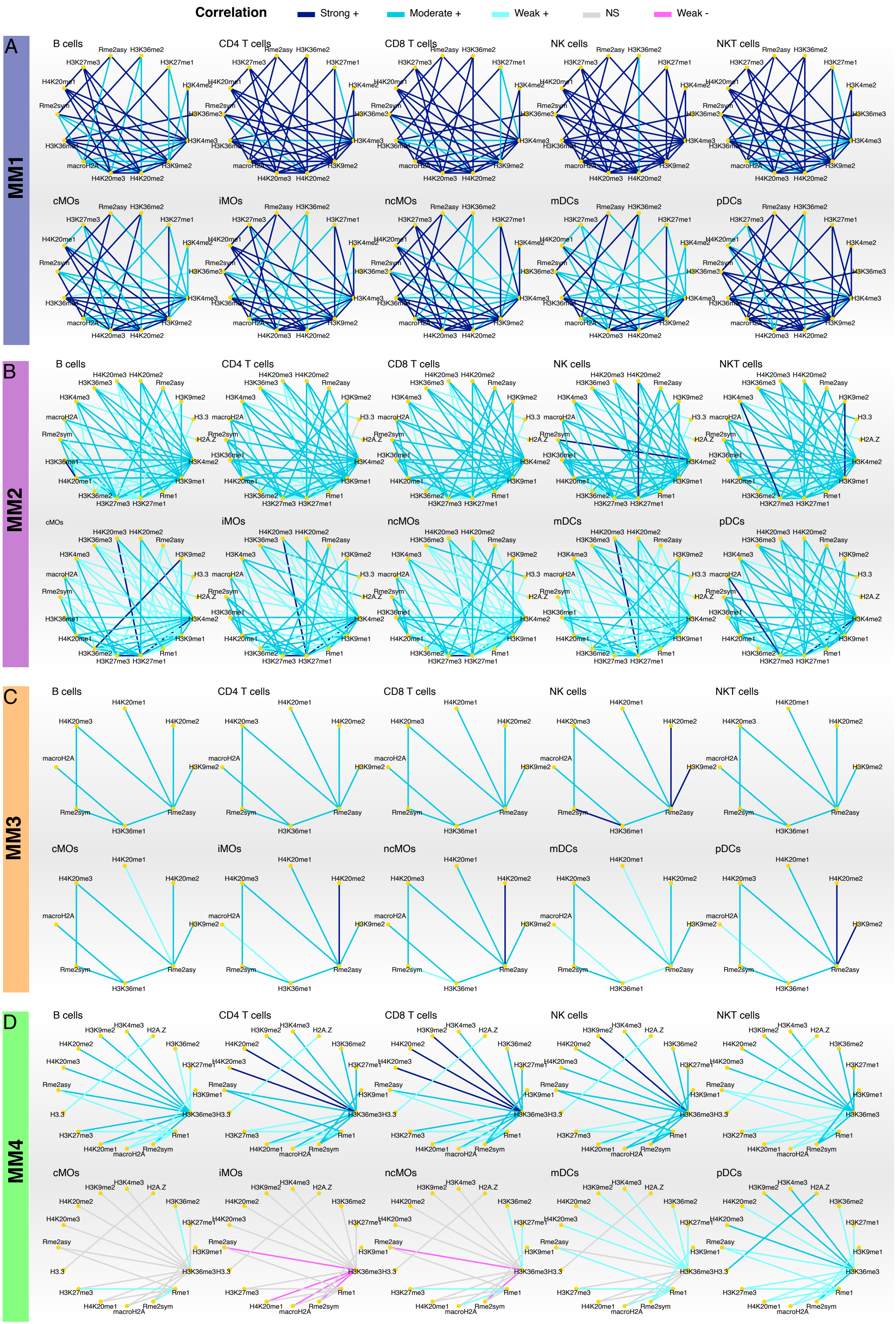


**Fig S2.** (Refers to Figure 3) Histone methylation modules in each immune cell type separately, except HPCs. **A** Methylation module 1. **B** Methylation module 2. **C** Methylation module 3. **D** Methylation module 4. The color of the edges represents stratified correlation coefficients.

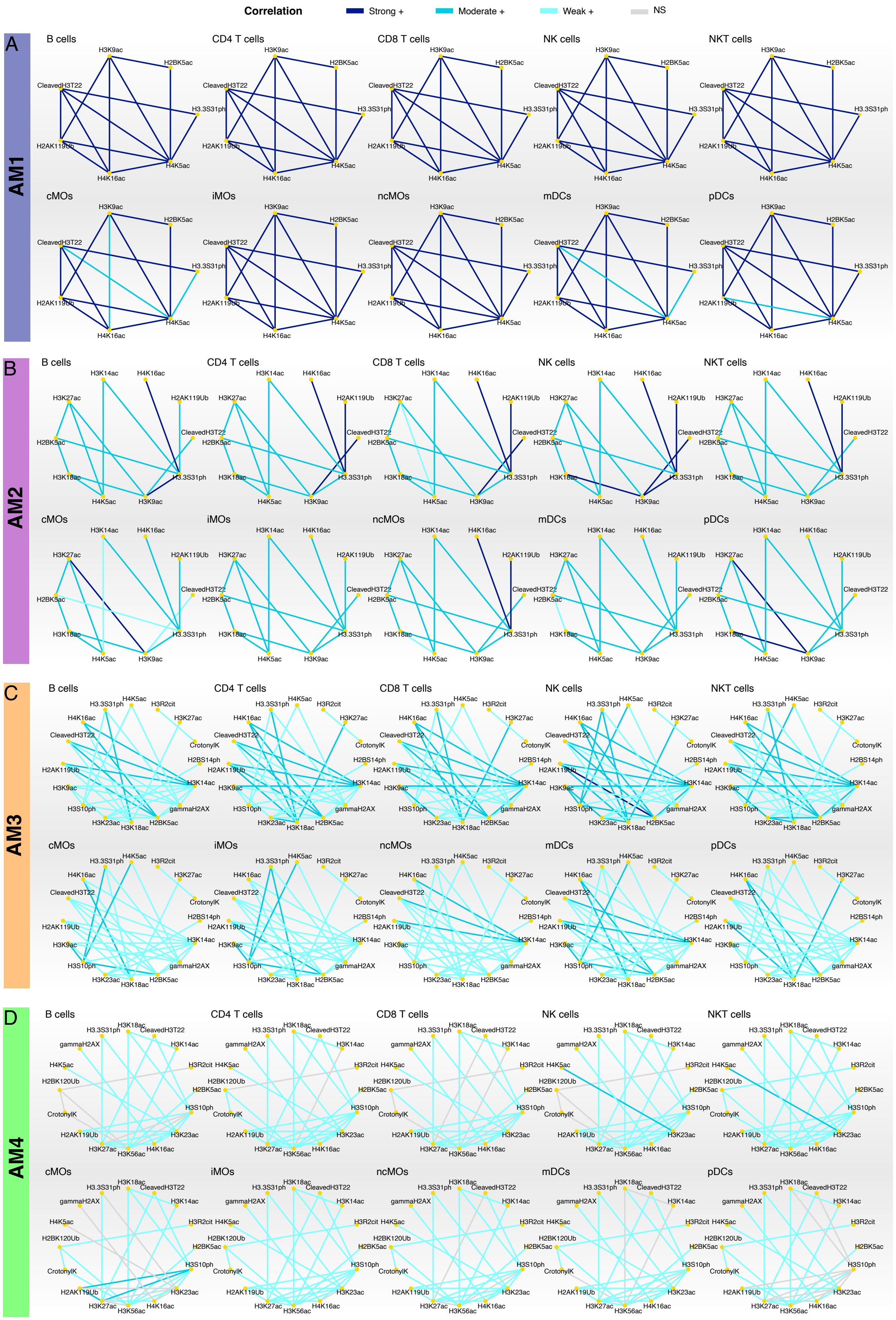


**Fig S3.** (Refers to Figure 4) Histone acetylation modules in each immune cell type separately, except HPCs. **A** Acetylation module 1. **B** Acetylation module 2. **C** Acetylation module 3. **D** Acetylation module 4. The color of the edges represents stratified correlation coefficients.

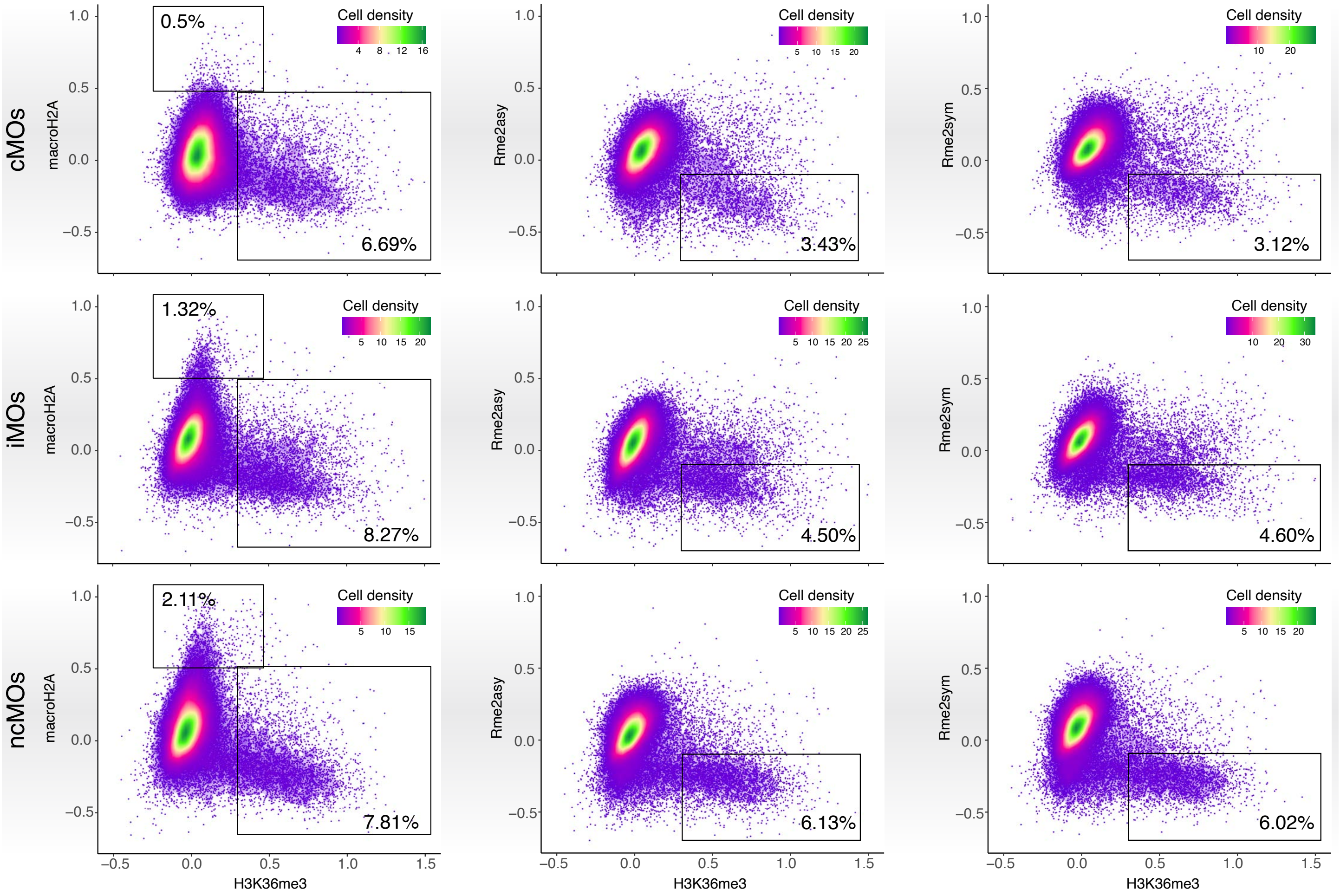


**Fig S4.** (Refers to Figure 3) Biaxial scatter plots of macro H2A, Rme2asy and Rme2sym against H3K36me3 showing proportions of different types of monocytes (cMOs, iMOs and ncMOs) that have increased abundance of H3K36me3. Scatter plots are rendered with cell densities (colored).

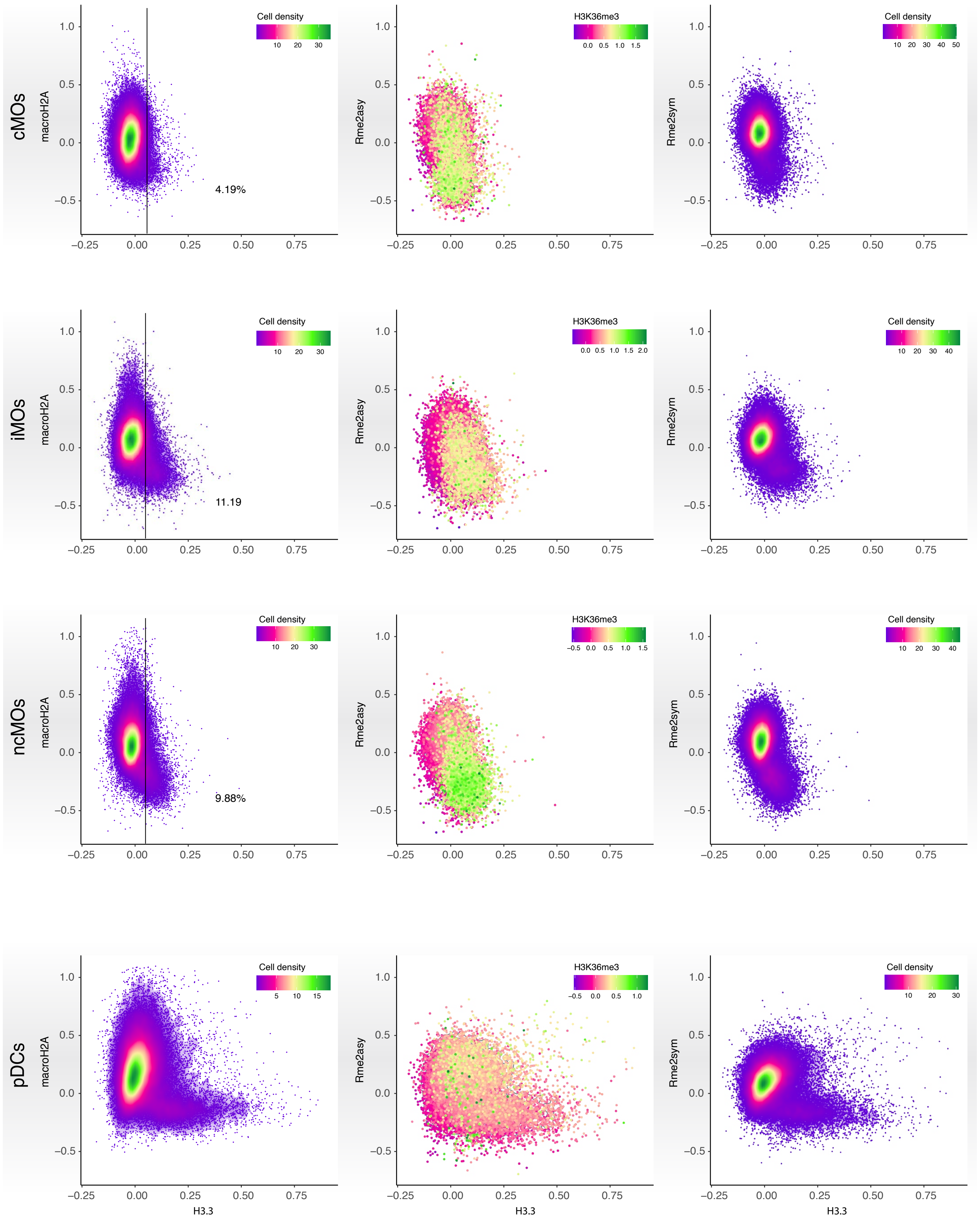


**Fig S5.** (Refers to Figure 3) Biaxial scatter plots of macro H2A, Rme2asy and Rme2sym against H3.3 showing different types of monocytes (cMOs, iMOs and ncMOs) and pDCs that have increased abundance of H3.3. Scatter plots are rendered with cell densities (colored, the first and third columns) or showing abundance of H3K36me3 (the second column). Increase of H3.3 coincided with the increase of H3K36me3.

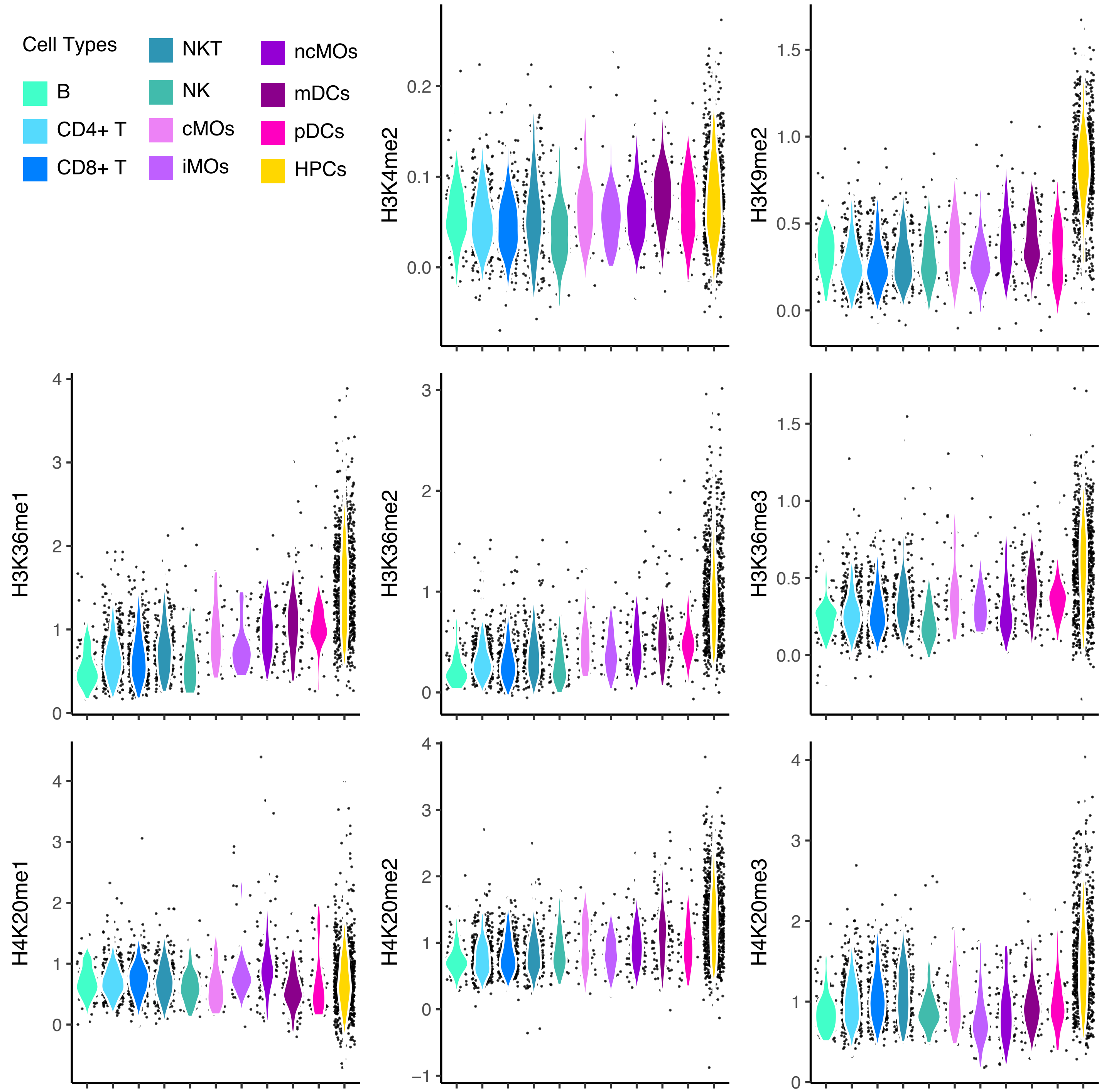


**Fig S6.** (Refers to Figure 5) Violin and point plots of all immune cells in MC6 showing abundances of selected HPTMs from the methylation panel. Progenitors (HPCs) contributed notably to the increased abundances of HPTMs.

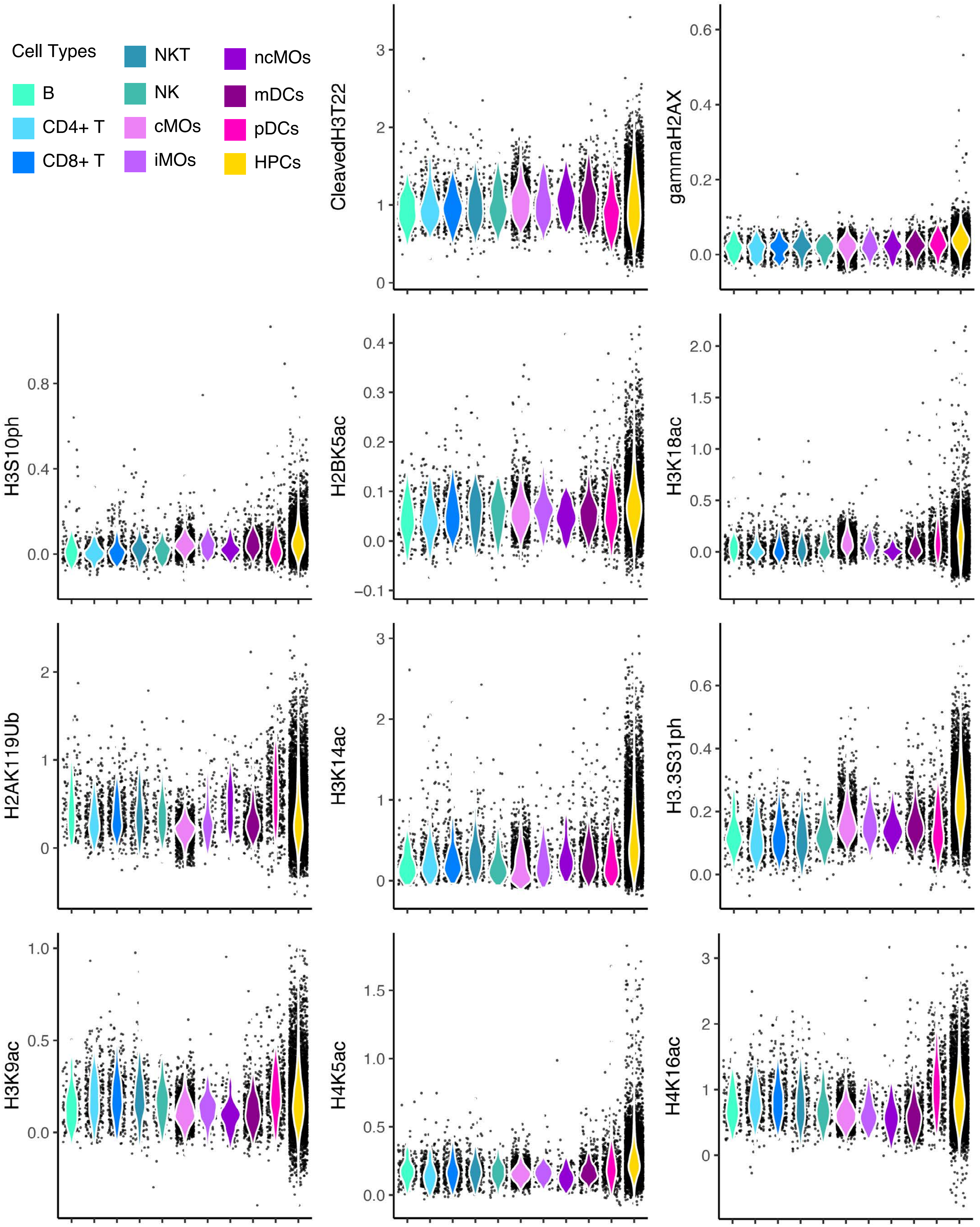


**Fig S7.** (Refers to Figure 5) Violin and point plots of all immune cells in AC5 showing abundances of selected HPTMs from acetylation panel. Progenitors (HPCs) contributed notably to the increased abundances of HPTMs.

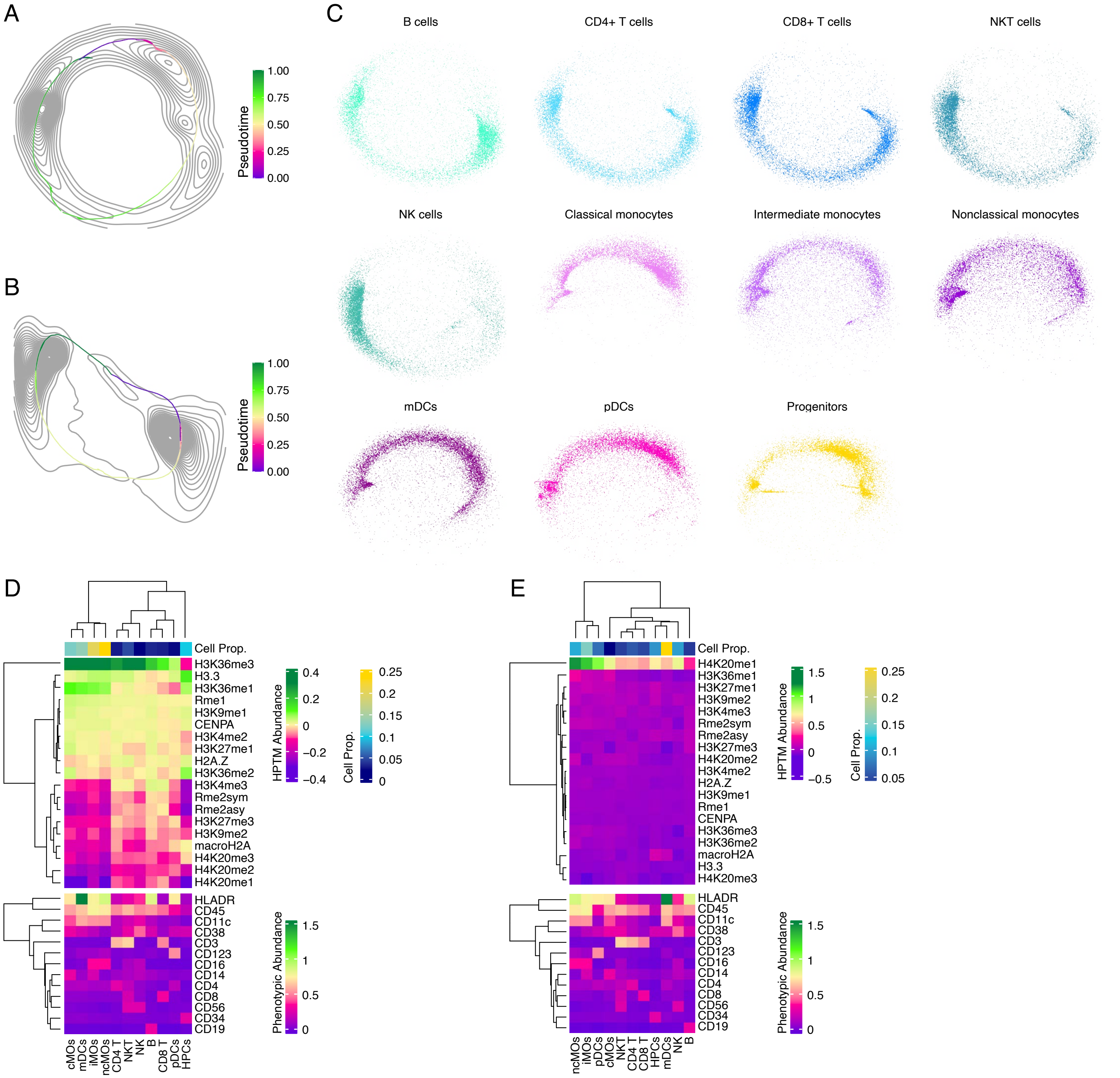


**Fig S8.** (Refers to Figure 6) Accompanying analyses of epigenetic trajectories. **A** Density plot of methylation trajectory with a fitted trajectory. **B** Density plot of acetylation trajectory with a fitted trajectory. **C** Methylation trajectory of each immune cell type shown separately. **D** A heatmap of Ex1 cluster shows HPTMs, phenotypic marks and proportion of immune cell types in this cluster. Cells of Ex1 have the highest abundance of H3K36me3. cMOs, iMOs, ncMOs, mDCs, and HPCs are the cell types with the highest proportions. **E** A heatmap of Ex2 cluster shows HPTMs, phenotypic marks and proportion of immune cell types in this cluster. Cells of Ex2 have the highest abundance of H4K20me1. mDCs, iMOs, and ncMOs are the cell types with the highest proportions. In **D** and **E** individual histone methylations are shown as z-score scaled values.
